# Movement is governed by rotational population dynamics in spinal motor networks

**DOI:** 10.1101/2021.08.31.458405

**Authors:** Henrik Lindén, Peter C. Petersen, Mikkel Vestergaard, Rune W. Berg

## Abstract

Although the generation of movements is a fundamental function of the nervous system, the underlying neural principles remain unclear. Since flexor- and extensor-muscles alternate during rhythmic movements like walking, it is often assumed that the responsible neural circuitry is similarly displaying alternating activity. Here, we present ensemble-recordings of neurons in the lumbar spinal cord that indicate that, rather than alternating, the population is performing a low-dimensional “rotation” in neural space, in which the neural activity is cycling through all phases continuously during the rhythmic behavior. The radius of rotation correlates with the intended muscle force and a perturbation of the low-dimensional trajectory can modify the motor behavior. Since existing models of spinal motor control offer an inadequate explanation of rotation, we propose a new theory of neural generation of movements from which this and other unresolved issues, such as speed regulation, force control, and multi-functionalism, are readily explained.

## Main

The neural circuitry behind movement encompasses several distinct forebrain regions, cerebellum and the brainstem. The core executive circuits for movement such as locomotion, however, reside in the spinal cord^1^. These spinal motor circuits, often referred to as central pattern generators (CPGs), are capable of autonomous generation of rhythmic coordination of muscles. Although great progress has been made in characterizing the cellular properties of spinal inter- and motor neurons, including their genetic lineages^2, 3^, the detailed network architecture as well as the associated neuronal ensemble dynamics remain elusive. Due to the apparent right-left and flexor-extensor alternation, it has often been proposed that distinct groups of interneurons, or ‘modules’, are active in a push-pull fashion and that the rhythm is ensured by cellular pacemaker properties^4, 5^. It is unknown if and how such organization and different motor programs are manifested in ensemble activity of spinal networks.

## Rotation in spinal motor circuits

Here, we examined the activity in spinal motor networks using extracellular multi-electrode recording in the turtle lumbar spinal cord. This preparation provided mechanical stability, which allowed simultaneous monitoring of large numbers of spinal interneurons in laminae VII-VIII and motoneurons during the execution of various rhythmic motor programs^6–8^. As expected, the activity of individual neurons was rhythmic in relation to the nerves, but the population activity as a whole appeared rather incomprehensible (**Fig. 1a-b**). However, when sorting these neurons according to phase of the motor nerve output we found that the population activity resembled a continuous sequence, that covered all phases of the cycle (**Fig. 1c**). To better understand the sequential activity, we performed a principal component analysis (PCA) of both the neuronal population and the nerve activity. Both the neuronal activity and the 6 motor nerves followed a low-dimensional manifold, i.e. most variance was explained by few components (**Fig. 1d**). Whereas the nerve activity appeared entangled, the neuronal activity had a simple rotation (**Fig. 1e-f**). Rotational population activity was independent on the sorting and it was observed in all trials and across animals (**Extended data Fig. 1-2, Supplementary Video 1**). To quantify this distinction further, we applied a previously defined metric^9^, that quantifies the “tangling” of neural trajectories, i.e. the degree to which points along the trajectory are close to each other, but move in different directions. We found the tangling to be larger for the muscle trajectories than the neuronal trajectories in the majority of the time (>96.3%), which was consistent across data sets (**Extended Data Fig. 3**). Since the tangling for rotational trajectories is lower than for trajectories with points that are close to each other and moving in opposite direction, as would be the case for alternating activity (**Extended data Fig. 1a**), these data are consistent with a neuronal population that is executing a rotation. There did not seem to be any discrete phase preference as otherwise expected in an alternating modular network (**Extended data Fig. 1-3**). Rotational dynamics has been observed in the motor cortex and elsewhere^10–12^, but it has not been described for spinal circuits previously. Nevertheless, indications can be found as wide phase-distributions in the scarce literature on spinal population recordings^6, 13–15^.

**Fig. 1.**
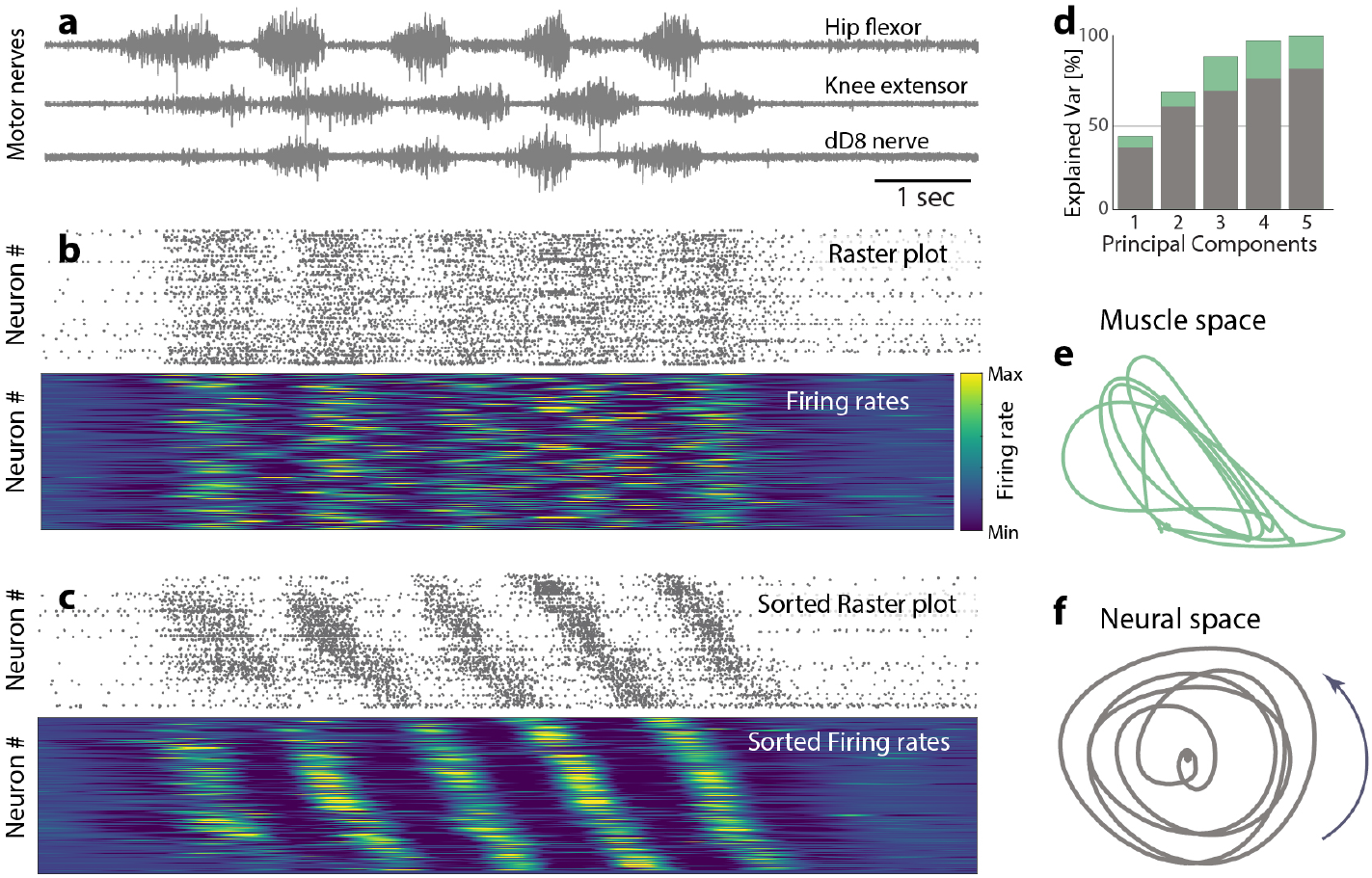
Neuronal population activity in the lumbar spinal cord has rotational dynamics. **a**, Activity of 3 selected motor nerves (ENG) during rhythmic hindlimb scratching movement. **b**, Concurrent ensemble activity in the turtle lumbar spinal cord as raster of spinal neurons (top, n=214) and estimated firing rates (bottom). **c**, Sorting the neurons in (**b**) according to phase (hip flexor) reveals sequential activity. **d**, First principal components explains most variance of ENG activity (green) and neuronal ensemble activity (gray). First two principal components of nerve activity, **e**, and neuronal population, **f**. Tangling of the nerve was higher than the network in 96% of the time.

## New theory to explain rotation and central pattern generation

Because conventional CPG theories, that are founded on a push-pull organization with intrinsically rhythmic modules^16, 17^, do not readily explain rotational dynamics, we sought to explore a new theory that can account for this and other open questions in spinal motor control. In particular, the mechanisms for generation of rhythms have remained nebulous. Cellular pacemaker properties has been suggested^4^, but decades of research has not been able to pinpoint a responsible cell type^17^. Here, we propose that the rhythm arises as a network oscillation rather than via cellular properties. It is well known that a network which is close to the transition point of dynamical instability, can have rhythmogenic properties without requiring specific cellular properties^18^. Since the CPG network structure is unknown, we parsimoniously assumed a structure, where glutamatergic neurons were randomly and recurrently connected. To prevent catastrophic run-away activity^19, 20^ the excitation (E) was balanced by recurrent glycinergic inhibition (I) (**Fig. 2a-b**), in line with reports of balanced synaptic input in various motor circuits^21–23^. Balanced networks of this type are known to undergo a phase transition when synaptic weights are increased beyond a critical value^24, 25^. For large networks, activity in this regime is chaotic^26^, whereas finite-sized networks in a dynamical regime close to the transition point may display more regular activity^27^. A linearization of the dynamics close to this point (see Mathematical Note, Supplementary information) demonstrates that finitely-sized networks can generate oscillatory activity if the leading eigenvalue of the connectivity matrix has a non-zero imaginary part^27^. Based on this idea, we set up a model network of rate-neurons with sparse connectivity where an external input, in the form of a synaptic drive (e.g. sensory-related or descending from the brain), could move the eigenvalues of the connectivity matrix across the stability line due to change in set-point of the firing-rate function (**Fig. 2c-d**). A second type of input that modulates the gain of individual neurons28 was also included to provide a mechanism to modify the network state. As the network received a sustained synaptic input, some of the eigenvalues moved beyond a critical level (red dashed line) which caused firing rates in the network to display self-sustained rhythmic activity (**Fig 2d-e**). When sorting the neurons according to phase, a sequential activity was revealed, i.e. a rotation, similar to the experimental observations (**Fig. 2f-g**). We refer to a network in this state as a balanced sequence generator (BSG). Both the BSG-model and the experimentally observed rotation are fundamentally different from conventional models, which are founded on alternation where the neurons have clustered phase preferences and belong to modules composed exclusively of either excitatory or inhibitory neurons.

**Fig. 2.**
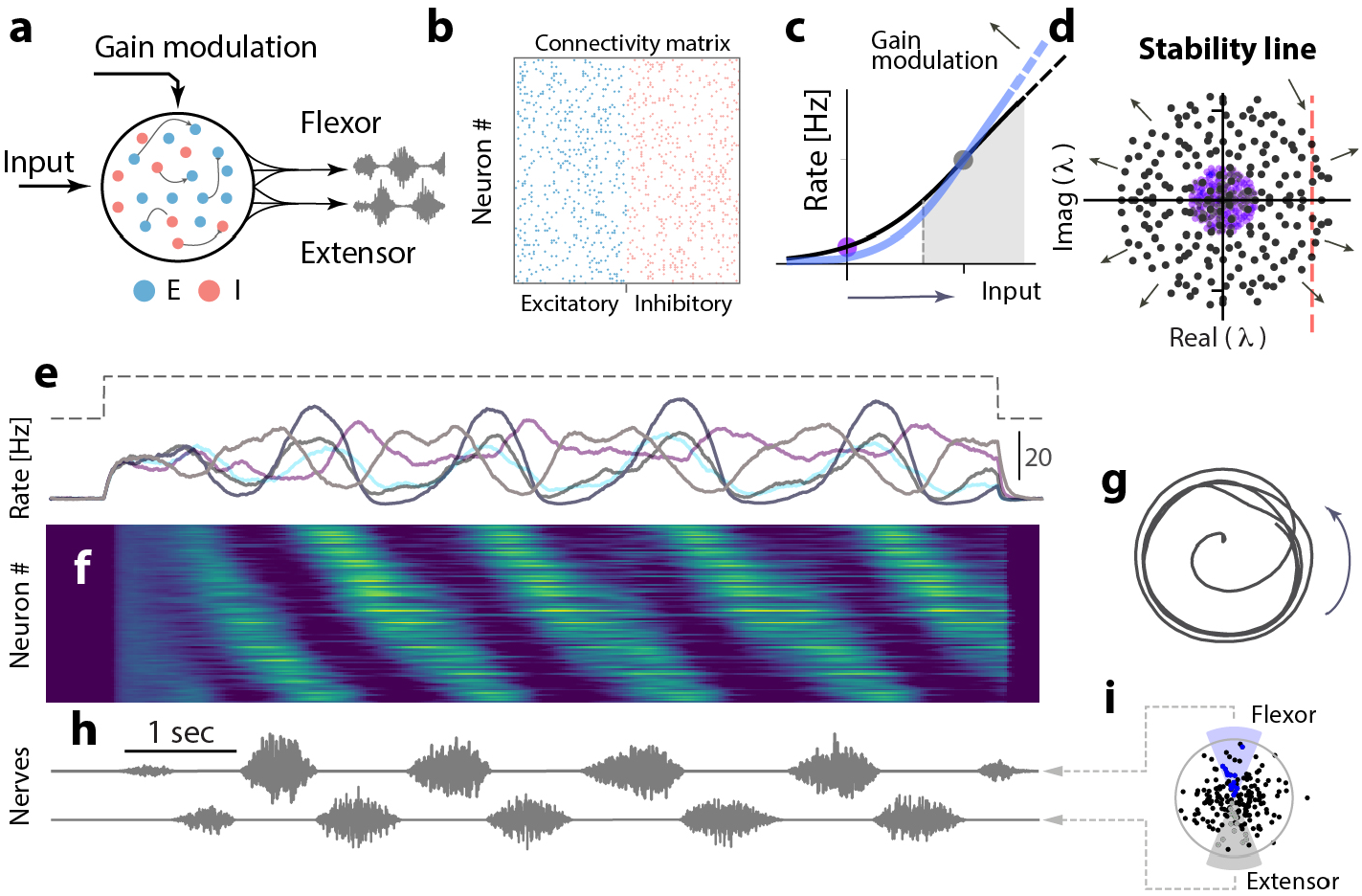
Rotational dynamics emerges in the BSG-model. **a**, Sketch of the BSG-model: An input drive activates a recurrent network with excitatory (blue) and inhibitory (red) neurons. The network can both receive synaptic input and gain-modulation. A subset of cells provides motor output. **b**, Connectivity matrix has 50% excitation and inhibition. **c**, The firing rate is increased by synaptic input (bottom arrow). This causes the eigenvalue spectrum to expand (purple vs. gray, **d**) and cross the stability line (broken red line) and thus generate a network oscillation. A gain modulation results in a change in slope (blue line and arrow, c). **e**, Input (top) and firing rates of 5 neurons (bottom). **f**, Sequential activity revealed by sorting according to phase, similar to experiments. **g**, Projection of the population firing rates on the two first PCs reveals a rotation. **h**, Model nerve output displays alternating activity. **i** Flexor/extensor nerves are innervated by anti-phase excitatory neurons in the strongest eigenmode (blue/gray).

To model the output nerve activity from the BSG-model we connected a subset of neurons based on their phase in the dominant eigenmode to pools of motor neurons to provide the appropriate nerve activity. This resulted in an alternating nerve output in resemblance with experiments (**Fig. 2h-i, Supplementary Video 2**). Next, we investigated the activity of the excitatory and inhibitory populations during the motor program in the BSG-model. It turned out that both the E- and I-populations themselves display similar sequences as the combined population activity (**Extended data Fig. 4**). These results demonstrate that rotational dynamics can arise in simple networks without fine-tuning of parameters and result in an alternating nerve output, in line with our experimental findings (**Supplementary Video 2**). Although proprioceptive feedback from muscles and their reflexive circuitry were not included in the BSG-model, we expect these to improve the performance by stabilizing the rhythmic activity.

## Control of force and period

Next, we evaluated whether the BSG-model could explain previously unsolved issues, such as independent control of force and speed of the movement. The ability to modulate the strength of the output and speed is key for volitional control but, to our knowledge, no mechanisms has been proposed for controlling these independently. To investigate these aspects in the model, we used gain modulation, i.e. the slope, (**Fig. 2c**) of the neuronal firing-rate function around the working point set by the external input28. First, we found that collective (uniform) modulation of the gain by an input drive could indeed control the amplitude in the BSG-model (**Fig. 3a-c**). As the amplitude increased so did the radius of rotation, while the frequency and sequence remained largely unaltered (**Extended data fig. 5**). To verify this prediction experimentally we inspected trials that, due to an inherent variability, had various radii of rotation (**Fig. 3e, Extended data figure 1c**). The radius of rotation had substantial correlation with the motor nerve activity (**Fig. 3f-g, Extended data figure 6**), in line with the predictions from the BSG-model and the proposed mechanism for amplitude control.

**Fig. 3.**
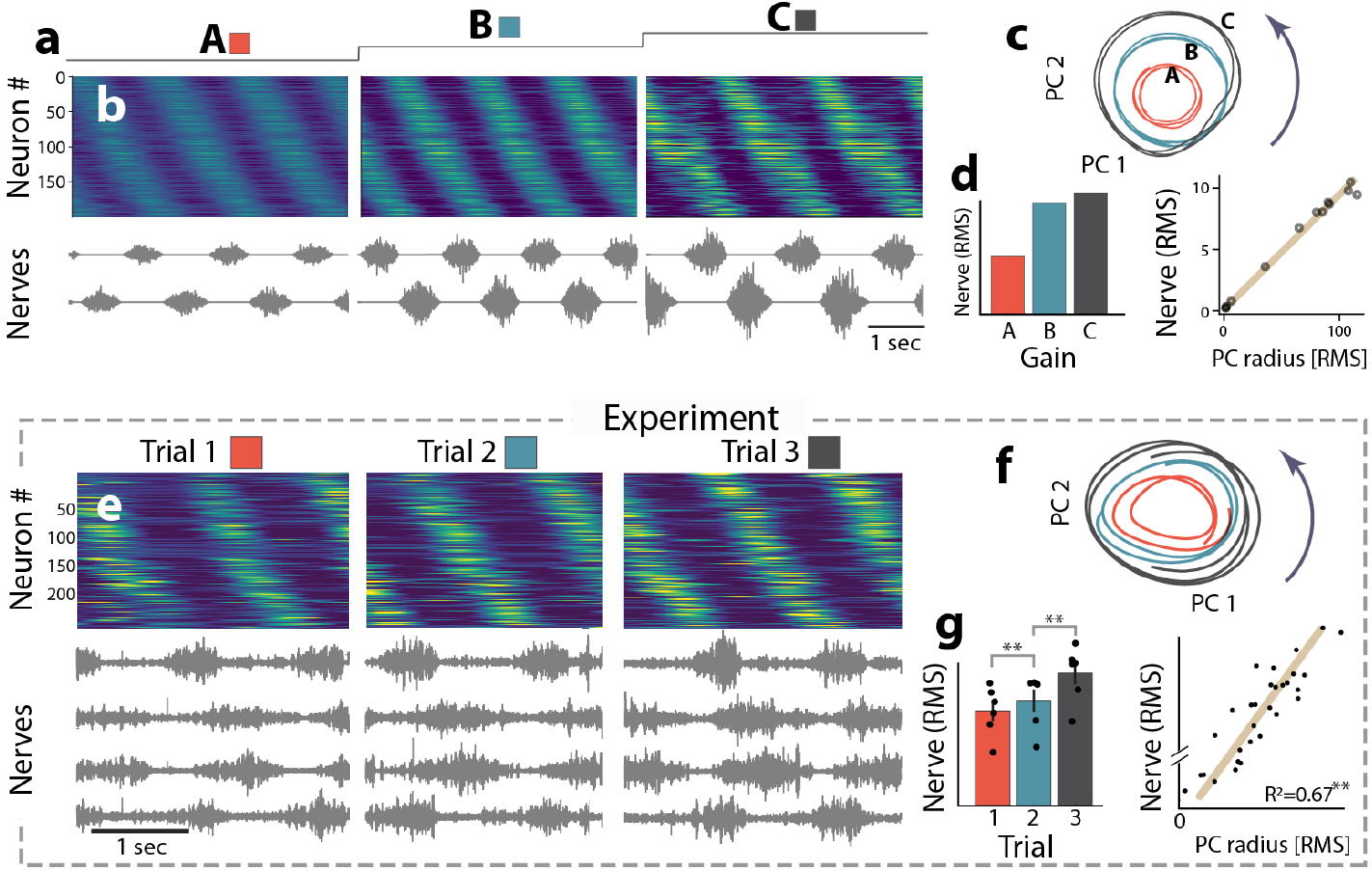
Network control of amplitude in the BSG-model. **a**, Increasing the common gain in the network (A–C) ramps up ensemble activity, **b**, and nerve output (bottom). **c**, Higher firing rates are associated with larger radius of rotation in PC-space (color-matched with levels A–C). **d**, Correlation between radius, gain and nerve output. **e**, Experimental verification: Trials 1, 2 and 3 with different nerve output and radius (**f**, color-matched). **g**, nerve amplitude (RMS±SE of 6 nerves) vs. radius of rotation (RMS of PC1 and 2). g, Wilcoxon test, Linear regression, **: *p* < 0.01, F-statistics.

Next, we explored if the BSG-model could control the period of the rhythm and thereby the speed of movement execution. Rather than collectively adjusting the neuronal gain in the network, we found that selective gain-modulation could alter the frequency of the population activity without affecting the amplitude (**Fig. 4**). Individual gain-modulation is a powerful tool in network control28 and here we systematically tuned the neuronal gain to identify a subset of neurons that had most influence on the period (**Fig. 4a-f**). Some neurons had a strong either positive or negative effect, which we call “brake-” and “speed cells”, respectively, while others had small effects on the rhythm. There were both inhibitory and excitatory neurons among both the speed- and the brake cells (**Fig. 4g-h**). Interestingly, cells with a speed-modulating capacity have been demonstrated experimentally^29, 30^. However, since both excitatory and inhibitory neurons were found among the brake- and speed cell categories in our model, an experimentally testable predication would be that also inhibitory neurons can have similar speed-modulating effects. The modulation capacity of individual neurons in the model is not due to their cellular properties, but rather their specific location in the network structure. A possible link between the network location, cell identity, and speed control remains to be assessed.

**Fig. 4.**
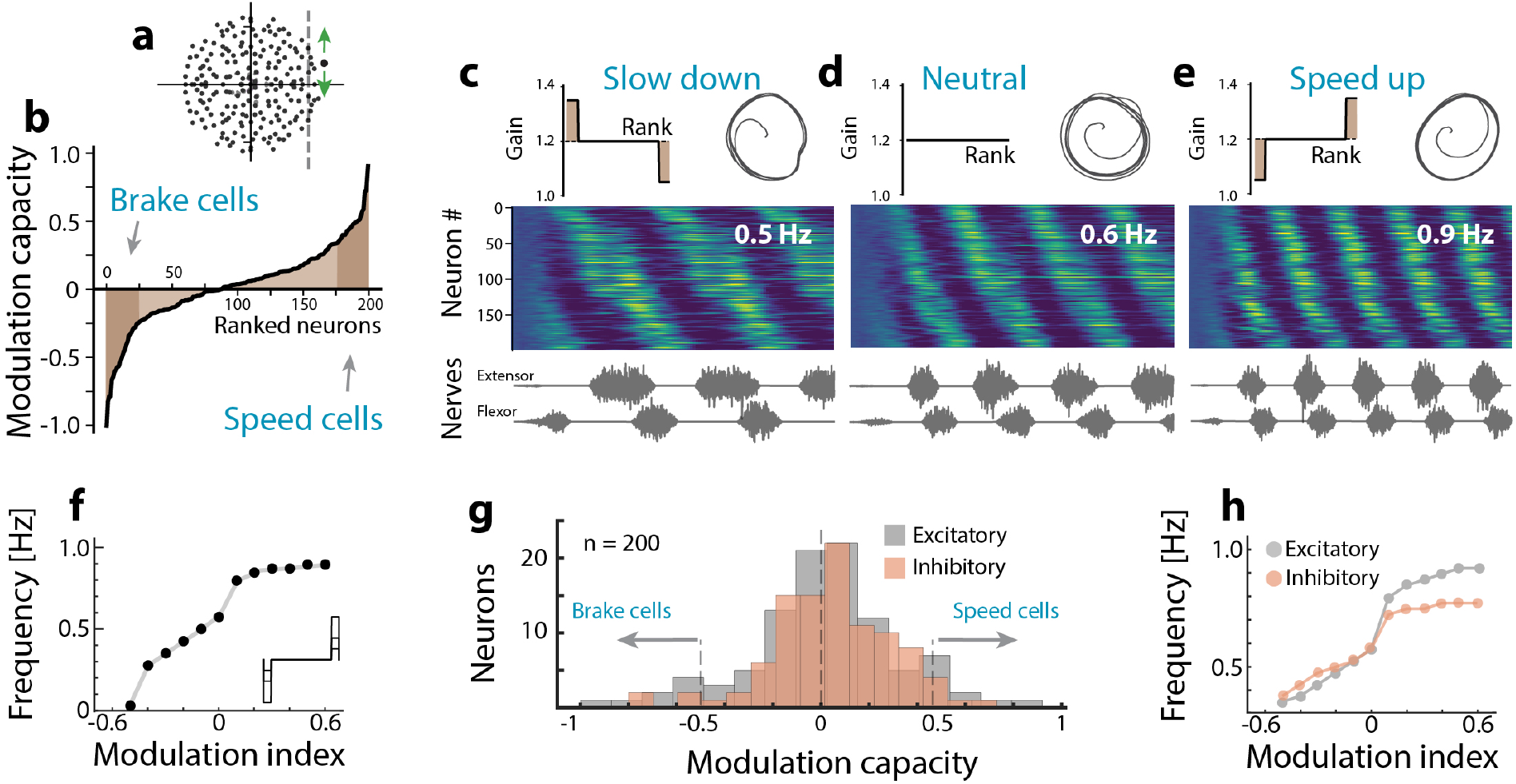
Modulation of period in the BSG-model. **a**, Adjusting the the neuronal gain neuron the rhythm by moving the eigenvalue up or down (green arrows). **b**, Capacity to modulate the rhythm by an individual neuron is assessed by changing its gain. Ranking neurons accordingly reveals “brake-” and “speed cells”. **c**, When activating brake cells while impeding speed cells (gain profile, top left), the rhythm is slowed down compared with neutral, **d**. Sorted ensemble activity (middle) and nerve output (bottom). Radius of rotation is largely unchanged (PCs top right) indicating a similar amplitude of motor output. **e**, Reversed activation results in faster rhythm (0.9 Hz). **f**, Gradually modulating the speed/brake cells (inset) can either decrease or increase frequency. **g**, Capacity to modulate the rhythm has a bell-shaped distribution. Brake and speed cells represent cells with strong modulation capacity, in which, both excitatory and inhibitory cells are found. **h**, Modulating only excitatory (gray) or inhibitory (orange) cells is sufficient to change the frequency.

## Population activity of multiple motor programs

The ability to execute multiple motor behaviors, i.e. a multifunctional output, is the hallmark of the motor system^31, 32^. Although cortical network models have already been demonstrated to generate multifunctional output^33, 34^, contriving a model within the conventional framework of spinal motor circuits that can accommodate the rich repertoire of behaviors has so far been a major challenge. Here, we focused on two well-known motor behaviors in the turtle and investigated these both experimentally and in the model. These behaviors consist of hind limb movements, where either the knee is protracted while moving the foot in small circles (pocket scratching) or protracting the foot with a stretched leg (rostral scratching)^35^. We hypothesized that this multi-functional activity is caused by a perturbation of the rotational dynamics that in turn switches the phases of the resulting motor nerve outputs. To test this idea in the BSG-model we identified two subsets of neurons for which two distinct sets of gain-modulation (“gain-profiles”, **Fig. 5a-b, Supplementary Video 3**) caused a moderate change in the phase preference of individual neurons. A comparison of the resulting neuronal phase-preferences between the two behaviors indicated that many of the neurons kept their timing in the sequence (**Fig. 5f-g**). We then optimized a set of read-out weights to drive motor nerve activity that caused a phase-shift of the hip angle between the two behaviors (**Fig. 5c**). In the resulting simulation the nerve output of behavior 1 had knee/hip extensors in phase (‘no shift’, **Fig. 5d**), whereas the second input pattern caused the phase of the hip extensor (and flexor) to change in relation to the knee extensor (**Fig. 5e**). Despite the marginal visual differences in population activity between the two behaviors (cf. **Fig 5d-e, Supplementary Video 3**), the network generated markedly different motor outputs. Using PCA, we found that the switch between behaviors was associated with a change of the low-dimensional subspace of the rotational dynamics. When projecting the population activity of behavior 2 onto the principal components (PCs) for behavior 1 (red) the rotational dynamics had a smaller variance compared to the variance of behavior 1 (black) (**Fig. 5h**). However, a comparison with the variance of the projection of the “native” PCs of behavior 2 (not shown) showed that this was not due to a markedly lower variance of behavior 2 compared to behavior 1, but instead that a fraction of the variance was in another subspace. By computing the ratio between variance explained in these two subspaces^36^ we quantified the subspace overlap between the two behaviors to be 0.49. These model results were qualitatively similar to the experimental data, where the sequential activation, although not identical, remained during the two behaviors (**Fig. 5i-m**). The subspace overlap here was 0.34 (**Fig. 5m**). Similar trend was seen across trials, behaviors and data sets (**Extended Data Fig. 7**). Finally, we tested whether other distinct motor patterns could be evoked in the BSG-model. A plethora of patterns or “gaits” could be induced via different gain profiles, with similar diversity to that of real motor patterns (**Extended Data Fig. 8-9**). This suggests that activating a spinal network to generate a desired motor pattern in general translates to finding the appropriate combination of neurons to modulate, e.g. by trial-and-error-based motor learning^37^.

**Fig. 5.**
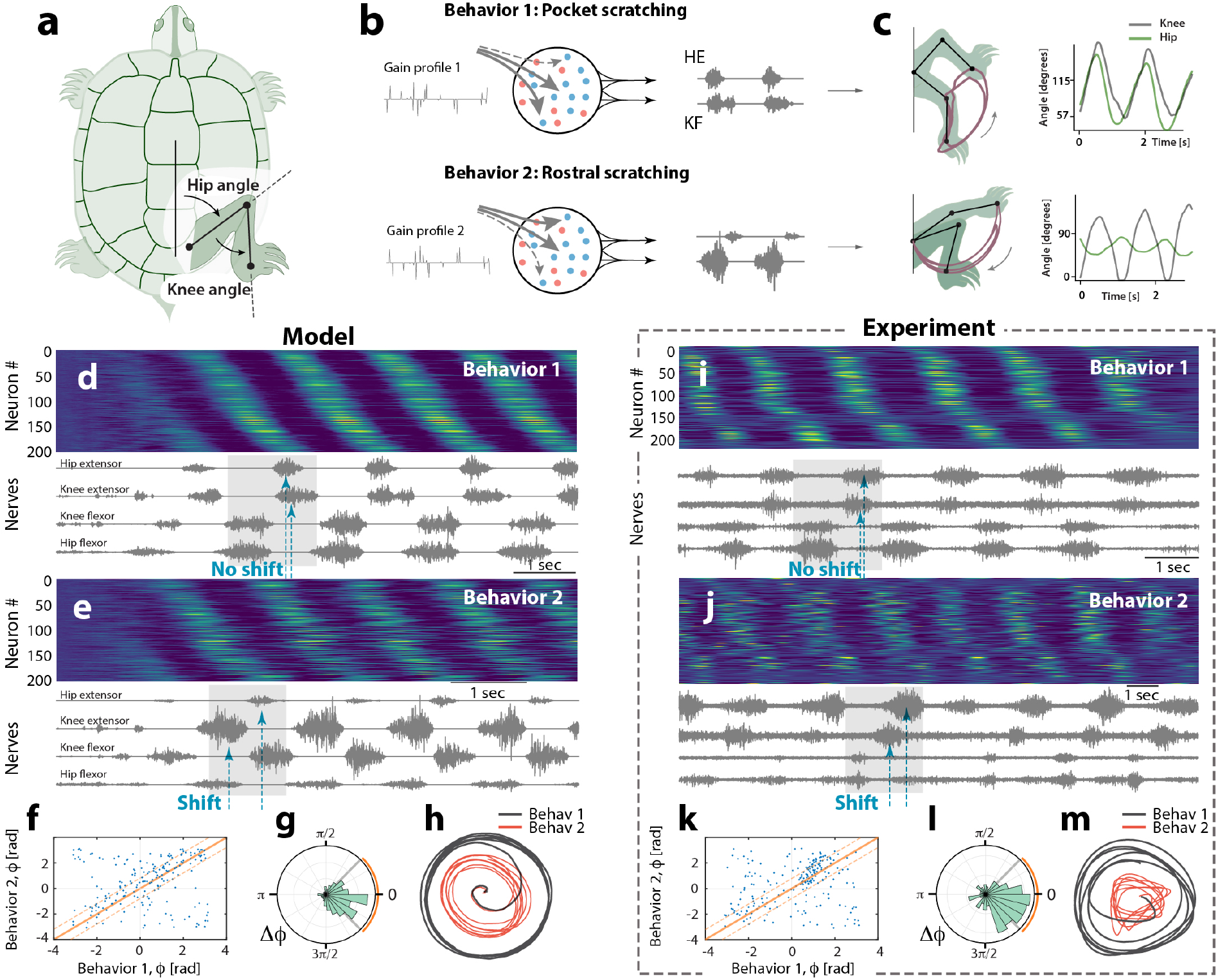
Multifunctionalism in model and experiment. **a**, hindlimb movement is quantified using the hip and knee angles. **b**, two distinct motor behaviors (pocket- and rostral scratching) are evoked in BSG-model by distinct gain profiles (left inset) of neuronal subsets. **c**, the evoked motor patterns (top: pocket, bottom: rostral) translated to limb trajectory (left, brown) and joint angles (right, hip and knee). **d-e**, associated ensemble activity are not identical, but have resemblance, although their motor patterns are qualitatively distinct (’no shift’ vs. ‘shift’ in shaded region). **f**, phase (*ϕ*) of neurons in behavior 1 vs. behavior 2 with respect to hip flexor scatter around the unity line, shown ±45°. **g**, phase difference (Δ*ϕ*) in polar histogram. Orange line indicates ±45°. **h**, Projection of behaviour 1 (black) and 2 (red) on the PCs of behaviour 1. Projection of behavior 2 (red) onto the subspace of behavior 1 had an overlap of 0.49 compared to its native representation^36^ using 3 PCs. **i-m**, experimental results in similar arrangement as (d-h) and similar motor behaviors. Projection of behavior 2 (red) onto the subspace of behavior 1 had an overlap of 0.34 compared to its native representation. (a) modified from35 with permission.

## Discussion

We have presented evidence that, rather than exhibiting alternating activity, the spinal network behind rhythmic movement displays low-dimensional dynamics that can be described as a rotation in neural space. During motor programs, the spinal population activity continuously cycles through all phases, while the resulting nerve activity is alternating (**Fig. 1**). Using computational modeling, we have shown that the core function of a spinal CPG, i.e. to convert a constant input to a rhythmic motor output, can be achieved by a simple balanced network that undergoes a transition to an oscillatory state. The alternating nerve activity is then obtained by a read-out from certain phases of the rotational population activity (**Fig. 2**). This model stands in contrast to conventional CPG theories that rely on cellular properties for rhythm generation and a modular hierarchy for pattern generation^4, 5^.

It is important to note, however, that our theory does not exclude the role of specific cell types^2^, e.g. for left-right coordination or speed control^29, 30^ and that cell-type specific connectivity could be included in the model to gain a theoretical understanding of its effect on resulting neural dynamics^38^. The BSG-model has all phases represented evenly in the population, which is a result of the simplified random connectivity (**Fig. 2b**). However, skewed phase representation could be achieved by including more structured connectivity, such as variable degrees of convergence and divergence while keeping the E/I balance. Experimental investigation of such architectures would require monitoring larger fractions of the network. Similarly, the role of intrinsic cellular properties, e.g. non-linear adaptation, could be included to elucidate their role in shaping network oscillations^27^.

This theory also offers an explanation of “deletions”, during which nerve burst are missing while the overall rhythmic pattern continues (**Extended data fig. 10**). A temporary disturbance of the rotational dynamics, which is large enough to bring the neural trajectory below the threshold for eliciting a nerve response, is sufficient to constitute a deletion (**Extended data fig. 11-13**). This suggests that a separation of spinal rhythm- and pattern-generating layers, as previously proposed^5^, is not necessary to account for deletions.

The ability to generate multiple movement patterns has already been studied for cortical networks^25, 28, 33, 34^, but the issue of multifunctionality in spinal motor networks has remained an open question. In our model, we explored a mechanism to generate multiple rhythmic motor patterns in the same spinal network by gain modulation of a subset of neurons in the network. Such subset modulation could be accomplished by cellular nodes that distribute sparse input to a larger population, as has been observed for spinal motor synergy encoders^39^.

Our theory could also be extended to account for non-rhythmic sequences by using a brief and targeted input drive, hence generating a single cycle of neural rotation, sculpted by selective gain modulation in the spinal network via descending commands from the brain. This could provide an important bridge between the motor circuits for rhythmic movement with those for non-rhythmic sequences, which is currently missing.

## Acknowledgements

Funding: This work was supported by The Independent research fund Denmark (no. 8020-00436B), and the Carlsberg foundation (no. CF18-0845). Mobilex. Lundbeck.

## Author contributions

R.W.B. conceived the original experiments. P.C.P. set up and performed the experiments, collected the data and analyzed some of the data. M.W., H.L. and R.W.B. conceived the original theory. H.L. performed the model simulations. H.L. and R.W.B. designed, and developed the theory, analyzed the experimental data, and wrote the manuscript.

## Additional Information

Supplementary Information is available for this paper. Correspondence and requests for materials should be addressed to RWB. Reprints and permissions information is available at www.nature.com/reprints.

