## Supplementary information for "Movement is governed by rotational population dynamics in spinal motor networks"

### METHODS

In this methods section, we describe the experimental protocols and the details of our computational modeling. The experimental data has been used in a previous study for a different purpose[1].

#### Experimental methods

The surgical procedures comply with Danish legislation and were approved by the controlling body under the Ministry of Justice (permission number 2018-15-0201-01504). Methods have previously been published in details[2–4]. Briefly, successful experiments on 4 adult red-eared turtles (Trachemys scripta elegans), ordered from Nasco (`https://www.enasco.com/`) of both sexes formed the basis of this study. One of the animals was used twice on different days, resulting in a total of 5 data sets. The animal was placed on crushed ice for 2 hours to ensure hypothermic anesthesia[3], then killed by decapitation and blood substituted by perfusion with a Ringer solution containing (in mM): 120 NaCl; 5 KCl; 15 NaHCO3; 2MgCl2; 3CaCl2; and 20 glucose, saturated with 98% O2 and 2% CO2 to obtain pH 7.6. The carapace containing the D4-S2 spinal cord segments was isolated by transverse cuts and the cord was perfused with Ringer's solution through the vertebral foramen, via a steel tube and silicone gasket pressing against the D4 vertebra. The motor nerves were cut to measure their activity and increase mechanical stability by preventing movements of the limbs. The preparation was placed and glued on the back in a chamber with constant flow of oxygenated Ringer's solution to keep the cord submerged and the skin tissue moist[3]. The vertebrae (D8-D10) corresponding to the lumbar segments L2-L5 in mammals[5] was carefully opened on the ventral side to allow access to spinal cord for insertion of the multi-electrode arrays. We opened the spinal column on the ventral side along segments D8-D10 and gently removed the dura mater with a fine scalpel and forceps. For each insertion site of the multi-electrode arrays, the pia mater was opened with longitudinal cuts along the spinal cord with the tip of a bend syringe needle tip (BD Microlance 3: 27G3/4", 0.4x 19 mm). The cuts were performed in parallel in the ventral horn between the ventral roots.

#### Electrophysiology

For monitoring the rhythmic activity and motor program state, electroneurogram recordings (ENG) were performed using suction electrodes of the hip flexor, knee extensor nerves and the dD8-nerve[6], i.e. a total of six motor nerves (3 from each side) at the level of D9-D10 vertebra. The ENGs were recorded with a differential amplifier (Iso-DAM8, World precision instruments, Sarasota, FL, USA) with filter bandwidth at 300Hz–1kHz. Custom–designed Silicon probes was inserted into the lumbar spinal cord (D8, D9 and D10) in the anterior-posterior direction to minimize damage to the white matter fiber tracks. These segments correspond to the lumbar (L2-L5) spinal cord in mammals[5]. Up to four 64-channel silicon probes, i.e. 256 recording sites, were inserted (Berg64 from NeuroNexus Inc., Ann Arbor, MI, USA). The probes had 8 shanks and 8 recording sites on each shank arranged in a staggered configuration with 30 $\mu$m vertical distance. The shanks thickness was 15$\mu$m and had a distance of 200 $\mu$m between shanks. Recordings were performed in parallel at 40 kHz using a 256 channel multiplexed amplifier (KJE-1001, Amplipex, Szeged, Hungary) to acquire the extracellular potentials of a large number of neurons, for post-hoc polytrode spike sorting.

#### Motor network activation by cutaneous sensory input

Each scratch episode lasted approximately 20 s. A new trial was initiated after a 5 min rest. To reproducibly activate the scratching motor pattern, a linear actuator was applied to provide mechanical touch on the skin around the legs meeting the carapace. The somatic touch was controlled by a function generator (TT2000, Thurlby Thandar instrument, UK) and consisted of a ten-second long sinusoidal movement (1-2 Hz). The touch was applied on the border of the carapace marginal shields M9-M10 and the soft tissue surrounding the hind limb, which is the receptive field for inducing pocket scratching motor pattern. Pocket scratching was elicited on either right or left side on the soft tissue surrounding the hind limb representing two distinct behaviors. Further, rostral scratching behavior was elicited by similar touching of the carapace in the more rostral location on the shields. For reviews on the various motor patterns and the cutaneous activation see[7,8].

#### Data analysis

All data analysis was performed in custom designed procedures either in Matlab (Mathworks, R2020b) or Python (www.python.org). Spike sorting was performed using KlustaKwik[9]. Spike rates were estimated by convolving the the neuronal spike times with a

Gaussian kernel,

$$k(t) = \frac{1}{\sqrt{2\pi}\sigma} \exp\left\{ \left( -\frac{t^2}{2\sigma^2} \right) \right\} \tag{1}$$

where $\sigma = 250$ ms, to capture the slow firing rate modulation. The firing rates were further high–pass filtered with a 3-pole Butterworth filter using a zero–phase filter ('filtfilt.m') function in Matlab, with a cut–off frequency of 0.3 Hz. To better visualize the phase of neuronal activity, the firing rate amplitude was normalized to unity and the mean subtracted. Further, to analyze the oscillatory part of the data, the slow components was removed in of the firing rates and nerve traces by digitally filtered using a 3–pole Butterworth filter in both directions to cancel phase distortion. The fast activity was high pass filtered with cut off 5 Hz after removing any potential action potentials. The slow activity was band–pass filtered from 0.2–5Hz. This data filtering was performed on all the data except the data involved in linear decoding, where the slow components has an important element in the translation between population spiking and the nerve output.

**Principal component analysis** (PCA) of the multidimensional population firing rates was performed on the firing rate space (neural space). The principal components $U_n$ were determined as eigenvectors of the empirical covariance matrix $C$ of the $n$ firing rate traces, with the eigenvalues $\lambda_n$ representing the absolute amount of variance in the data in that each component can account for. The eigenvectors and eigenvalues were found via:

$$\boldsymbol{CU} = \boldsymbol{U\Sigma} \tag{2}$$

where $\boldsymbol{U} = [U_1, U_2, ..., U_n]$ contains the principal components (eigenvectors) $U_n$ and

$$\boldsymbol{\Sigma} = \begin{bmatrix} \lambda_1 & 0 & \dots \\ \vdots & \ddots & \\ 0 & & \lambda_n \end{bmatrix} \tag{3}$$

The PCA was performed in matlab using the function 'PCA.m'. Similar PC analysis was performed on the nerve activity, although this was only 6-dimensional data (Fig. 1). The neuronal population activity plotted in PC-space as a function of time was achieved by projecting the population vector, $\vec{r}(t) = [r_1(t), \dots, r_n(t)]$, i.e. a vector with the firing rates of all neurons, onto the principal components, giving the population vector in new coordinates, $\vec{r}'(t)$:

$$\vec{r}'(t) = \vec{r}(t)\boldsymbol{U} \tag{4}$$

**Subspace overlap analysis** A method for quantifying to which extent a principal component subspace of one behavior overlaps that of a different behavior has been introduced previously[10]. To quantify the overlap between the low-dimensional subspaces of two different behaviours we first computed the principal components (PCs) of the two behaviours separately using PCA (see above). We then selected the first three PCs as their respective subspace. The overlap between subspaces was then calculated as the total variance captured by a projection of the first behaviour on the PCs of the other behavior, divided by the variance captured by the projection onto its "native" PCs[10]. We used the first 3 PC dimensions for this quantification in both experiments and model simulations (Fig. 5 and **Ext. Data Fig. 7**).

**Sorting of units according motor phase** The firing rate of units was sorted according motor phase by two steps. First, the frequency of rhythmic activity was identified by estimating the peak in the power spectrum of a representative nerve. For this purpose, the nerve activity was rectified and smoothed and sub-sampled to have the same sampling rate as the estimated firing rates. Second, the magnitude and phase of the coherence $Coh_i$ between this nerve activity and the firing rate of the $i$th neuron was estimated via[11]:

$$Coh_i(f) = \frac{\sum_{j=1}^{k} R_{ij}(f) N_j(f)}{k\sqrt{S_{xxi}^2 S_{nerve}^2}} \tag{5}$$

where $k$ is the number of multi-taper spectral estimates ($k = 4$). $R_{ij}$ and $N_j$ are the individual taper spectral estimates using discrete Fourier transform of the firing rate of the $i$th neuron, $r_i(t)$, and the rectified and low-pass filtered nerve trace n(t):

$$R_{ij}(f) = \sum_{t=0}^{T} e^{j2\pi ft} r_i(t) w_j(t)$$

$$N_j(f) = \sum_{t=0}^{T} e^{j2\pi ft} n(t) w_j(t)$$

and $w_j(t)$ is the $j$th taper function, the discrete prolate spheroidal (Slepian) sequences[12]. These taper functions and the Fourier transforms were calculated using the matlab function 'dpss.m' and 'fft.m'. The power spectra of the firing rate of the $i$th neuron and the nerve were calculated as $S^2_{xxi} = \frac{1}{k}\sum_{j=1}^{j=k} R_{ij}R^*_{ij}$ and $S^2_{nerve} = \frac{1}{k}\sum_{j=1}^{j=k} N_j N^*_j$, where the $*$ indicate complex conjugate. The phase of the $i$th neuron was chosen from $Coh_i(f)$ at the frequency where the strongest peak in $S^2_{nerve}$ was found, which was the rhythm of the motor pattern. Based on the phase, the neurons was sorted and plotted (e.g. Fig. 1c).

Significance level could be assessed in coherence (although not used in this context) if the magnitude of coherence exceeded the following:

$$|Coh| > \sqrt{1 - p^{1/(nk-1)}} \tag{6}$$

where $p$ indicates the level of confidence, i.e. a 95%-confidence has $p = 0.05$. The degrees of freedom is $k$, and since we used $k = 4$ tapers for spectral estimation, the confidence limit was at 0.79. The standard deviation was calculated using circular statistics as originally defined in[13] (section 2.3.3):

$$\sigma_{circular} = \sqrt{-2log(\bar{R})} \tag{7}$$

i.e. involving the natural logarithm of $\bar{R}$ is the mean resultant length of all observations in polar coordinates, hence $\bar{R}$ is between 0 and 1. If the observations angles are close to each other the resultant length is close to 1 and the $\sigma_{circular}$ is close to 0.

**Nerve activity measures** In some of the analysis the motor output was measured as electroneurograms (ENGs) quantified using the root mean square (RMS) of the traces after smoothing using the Savitzky-Golay finite impulse response filter. The RMS is the square root of the mean of the squared ENGs:

$$ENG_{RMS} = \sqrt{\frac{1}{n}(x_1^2 + x_1^2 + \cdots + x_n^2)} \tag{8}$$

where $x_1, x_2, \ldots, x_n$ are the ENG measurements and $n$ is the number of samples. The RMS values were calculated in matlab using the procedure 'rms.m'. The mean values reported (Fig. 3g) are the average of all 6 nerves. The error bars are the standard error of the means, i.e. the standard deviation divided by $\sqrt{6}$. A pairwise statistical comparison was performed between trials each having 6 measurements (the nerves) using the non-parametric Wilcoxon signed rank test via the procedure 'singrank.m' in matlab.

The relationship between radius of PC rotation (RMS of first two components) and nerve output (RMS) was verified using an F-statistic vs. a constant model. Test statistic for the F-test on the regression model (**Extended Data Fig. 6g-h**), is a tests of whether the linear fit is significantly better than a constant.

**Nerve output prediction using a linear decoder** Linear decoding of neural ensembles e.g. in primary motor cortex has been used efficiently to control of prosthetic devices using a Brain-Computer Interface for individuals with tetraplegic conditions[14,15]. The idea is to use a linear filter, $\boldsymbol{f}$, i.e. a linear decoder, that can translate the firing rates of a population of neurons, written as a matrix $\boldsymbol{R} = \vec{r}(t)$ over a time period, to a read-out to control a set of muscles, $\boldsymbol{N}$, such that $\boldsymbol{N} = \boldsymbol{Rf}$. The filter is first constructed from training data that describes the association between firing rate matrix $\boldsymbol{R}$ and the the nerve output matrix $\boldsymbol{N}$ (**Ext. Data Fig. 12-13**). The filter was estimated using the least-squares formulation from a closed-form expression[16]:

$$\boldsymbol{f} = (\boldsymbol{R^T R})^{-1}\boldsymbol{R^T N} \tag{9}$$

In this study, we form a prediction of the nerve output based on the linear decoding of the neuronal population activity in the spinal cord, for the intention to verify how well such a simple measure can predict the output. In particular, this is relevant for the investigation of "deletions". The prediction of "deletions" purely from the sampled population activity can give insight to whether there are several layers in the motor network, i.e. separation of rhythm and pattern generation, which has previously been proposed for non-resetting deletions in decerebrated cats and neonatal isolated rodent spinal cords[17–20].

**Trajectory tangling metric of neuronal population and nerve activity** The degree of tangling of the trajectories in neural space compared with the motor nerve trajectories have recently become of interest[21,22]. We use the metric for trajectory tangling, $Q(t)$, previously defined[21] (**Extended Data Fig. 3**). The point in the multidimensional state space, $\dot{\mathbf{r}}(t)$, can either represent the population firing rate or the activity of the group of motor nerves (6 in our case), or the principal components thereof. The tangling is defined as the maximum squared Euclidean difference in velocity of the movement along the trajectory at two points in time, $t$ and $t'$, $\dot{\mathbf{r}}(t) - \dot{\mathbf{r}}(t')$, divided by the Eurclidean distance betweeen the points squared:

$$Q(t) = \max_{t'} \frac{\|\dot{\mathbf{r}}(t) - \dot{\mathbf{r}}(t')\|^2}{\|\mathbf{r}(t) - \mathbf{r}(t')\|^2 + \varepsilon} \tag{10}$$

This fraction is a basic measure of how different the velocity (speed and direction) is between two points on the curve divided by how far they are from each other. If the trajectory is very tangled, there will be points that have different directions and are close to each other. Parts of the trajectory with low tangling will tend to move in same direction, if they are close to each other. The $\varepsilon$ is a small constant added to avoid division by zero. The value of $\varepsilon$ is not important as long as it is small compared with the scale of the data. Similarly, the scale of the data should not affect the tangling metric as long as it is large compared with $\varepsilon$. Since we are comparing firing rates and ENG nerve recording, which are several orders of magnitude in difference, we scaled $\varepsilon$ by the root-mean-square of the first principal component. The derivative was estimated as the difference in $\mathbf{r}$ between neighboring samples and divided by the sampling time. Since this method tend to enhance noise, we first smoothed the trace with a kernel (500 point, Savitzky-Golay of 2nd degree). The unit of Q is seconds$^{-2}$. We calculated the tangling of the first 3 principal components of firing rates of the neuronal population and compared it with the tangling of the first 3 components of the 6-dimensional nerve activity. We used the fraction of time-points that are larger for the nerve activity than the network as a composite measure to compare across trails and animals (**Extended Data Fig. 3**).

### Network model: Balanced Sequence Generator

The model consists of a network of interneurons and two or more nerve-readouts that represents the motor commands resulting from the network activity.

##### Interneuron network

The interneuron network consists of $N = 200$ neurons out of which half are excitatory and half are inhibitory. We model the activity of an example neuron $i$ as a firing rate $r[g_i(t), V_i(t)]$ that depends on an activity variable $V_i(t)$, analogous to a membrane potential, and gain variable $g_i(t)$. We use a similar function to previously published models[23] adjusted avoid negative firing rates:

$$r(g_i, V_i) = \begin{cases} V_*(1 - \tanh[g(V - V_*)/V_*]), & \text{for } V \le V_* \\ V_* + V_{\max}\tanh[g(V - V_*)/V_{\max}], & \text{for } V > V_* \end{cases} \tag{11}$$

where $V_*$ represents the input level at which the slope of the firing rate function has its maximum (resulting in an output firing rate of $r = V_*$ Hz) and $V_{\max}$ is the maximum deviation (in terms of firing rates) from $V_*$. Here, we set $V_*$=20 and $V_{\max}$=50, resulting in a maximum firing rates of 70 Hz. The dynamics of the network is determined by the equation

$$\tau \dot{V}_i(t) = -V_i(t) + \sum_j W_{ij} r[g_j(t), V_j(t)] + I_e(t) \tag{12}$$

where $\tau$=50 ms is a time constant representing the combined membrane and synaptic timescale, $W$ is a matrix that describes the network connectivity (see below), and $I_e(t)$ is a time-varying external drive that consists of a constant input and a noise term $I_e(t) = I(t) + \varepsilon$, where the noise term $\varepsilon$ is Gaussian noise with zero mean and standard deviation $\sigma = 4$. The network thus receives two types of external input: a common external input ("drive") $I_e$ that is used to cause a transition from an inactive background state to an active rhythmic state (Fig. 2), and an input that sets the gain $g_i$ of individual neurons that is used to modulate the network activity in terms of amplitude (Fig. 3), frequency (Fig. 4) or for multi-functional behaviour (Fig. 5). For simplicity we used a constant input drive $I_e$=20 when studying the effects of gain modulation in the network.

##### Network connectivity

The connectivity of the network is assumed to be sparse (as indicated by e.g.[4]) with a pair-wise connection probability $C = 0.1$. Excitatory (positive) weights $w_{ex}$ are assumed to be equal in magnitude to inhibitory (negative) ones $w_{in}$. To ensure that the incoming connections are balanced for each neuron we construct the connectivity matrix $W$ in the following way: We start with a matrix where all elements are zero. For each neuron we then select $CN/2$ presynaptic excitatory neurons and assign them the weight $w_{ex}$ and $CN/2$ presynaptic inhibitory neurons and assign them the weight $w_{in}$. In this way we ensure that the network is both globally and locally balanced[24], i.e. the incoming synaptic weights are balanced for each neuron. The synaptic weight is set so that the connectivity matrix $W$ has a spectral radius (Fig. 2d) such that the largest eigenvalue $\lambda_{max}$=1 (on average over network realizations), by setting

$$w_{ex} = \frac{1}{\sqrt{NC(1-C)}} \tag{13}$$

and $w_{in} = -w_{ex}$[25]. This results in a network that is on the edge of instability for a uniform gain $g$=1. As a default, we set $g$ =1.2 which results in a linearly unstable network. For this study we selected connectivity matrices for which the largest eigenvalue $\lambda_{max}$ had a non-zero imaginary part since these networks can be expected to generate oscillatory activity (see Mathematical note below).

##### Gain modulation for amplitude control

To control the amplitude of oscillations in the network model we adjusted the gain parameter $g$ uniformly for all neurons in the network.

##### Gain modulation for frequency control: "Speed-" and "brake" cells

To control the frequency of oscillations in the network we adjusted the gain $g_i$ individually for selected neurons in the network. A simple procedure was set up to estimate the influence of each neuron on the overall frequency: The gain $g_i$ was increased and decreased by a small amount and the spectrum of the connectivity matrix $W$ was calculated (Fig. 4a). Depending on whether that imaginary part of the largest eigenvalue $\lambda_{max}$ was increased or decreased (corresponding to an expected increased or decreased oscillation frequency) we assigned the neuron an index depending on its frequency "modulation capacity". A positive modulation capacity means that an increase in gain or drive to that neuron will increase the frequency of the rhythm, and vice versa for a negative modulation index. Since a detailed gain modulation of all neurons in the network can be considered less biologically plausible, we selected the 10% of neurons with the largest positive effect on the imaginary part and labelled them as 'speed' cells, and the 10% with the largest negative effect and labelled them 'brake' cells. To increase the network oscillation frequency we increased the gain of the 'speed' cells and decreased the gain of the 'brake' cells (Fig. 4). To decrease the network oscillation frequency we did the opposite, i.e. we decreased the gain of the 'speed' cells and increased the gain of the 'brake' cells.

##### Gain modulation for multi-functional activity: "Switch" cells

To generate different motor behaviour from the network we identified a subset of neurons that had a large influence on the phase distribution of the dominant eigenmode. Starting with a default value for the gain of $g$ =1.1 we first calculated the phase for each interneuron from the eigenvector corresponding to the largest eigenvalue of the connectivity matrix $gW$. We then increased the gain $g_i$ of each neuron $i$ individually and calculated the effect of the changed gain on the phase distribution of the resulting effective connectivity. The top 10% of the neurons that caused the largest change in the overall phase distribution (calculated as the circular standard deviation of the change in phase) were selected as 'switch' neurons. To generate two different distinct behaviours, we set the gain of the 'switch' neurons to two different random vectors with values uniformly distributed between $g_i = 1.1 \pm 0.3$.

##### Nerve readout

The nerve activity was modeled using a Gaussian noise with zero mean and where the width (standard deviation) $\sigma(t)$ of the distribution depends on a threshold-linear readout from the interneuron network:

$$\sigma(t) = [\sum_i M_i \phi_i(t)]_+ \tag{14}$$

where $M_i$ represents the readout weights and $[]_+$ indicates that the width can only be positive. The readout weights were constrained to respect Dale's law, i.e. excitatory interneurons could only have a positive weights and inhibitory interneurons could only have negative weights. We used two different ways of setting up the linear readout:

**Readout based on phase of dominant eigenmode** The simplest method used was to set up to readout-weights $M_i$ based on the phase of each neuron $i$ in the network oscillation. The phase of all neurons was estimated from the eigenvector corresponding to the largest eigenvalue $\lambda_{max}$ of the connectivity matrix $W$. To set up the readout for a specific nerve, we first assigned the nerve a phase $\theta_{nerve}$. For excitatory neurons that had a phase of $\theta_{nerve} \pm \pi/8$ we set $M_i = 1$ and set $M_i = 0$ for all other excitatory neurons. To generate reciprocal inhibition in the nerve input, we selected inhibitory neuron with a phase of $(\theta_{nerve} + \pi) \pm \pi/8$ and set $M_i = -1$, while $M_i = 0$ for all other. To set up a pair of flexor-extensor nerves with alternating activity, we set $\theta_{flexor} = \pi/2$ and $\theta_{extensor} = -\pi/2$.

**Optimized readout for multi-functional output** We first selected two distinct gain vectors for pocket and rostral scratch, respectively (see above) and simulated network activity using these gain vectors. To find the appropriate read-out weights, we then set up sinusoidal target functions for the nerve function "input" (i.e. the sum in eq.14) for each behaviour and for each nerve separately. The flexor- and extensor nerves were phase-shifted by $\pi$. The pocket and rostral scratch behaviours had different relative timing between the knee- and hip flexor nerves, shifted by $\pi$ as well as different amplitudes (Fig 5). Read-out weights were then found using a linear least-square algorithm with bounds on the variables (implemented in Python using `scipy.optimize.lsq_linear`) such that the weights $M_i$ could only be positive for excitatory neurons and negative for inhibitory neurons.

**Limb movement model** To translate the nerve readout to position of knee and foot we set up a simple model that integrates the nerve drive to calculate the angle $\Theta$ of the foot/knee joint resulting from the flexor and extensor nerves (Fig. 5 and Supplementary Video 3):

$$\tau_{muscle}\dot{\Theta}(t) = w_{\Theta}[flexor(t) - extensor(t) - (\Theta - \Theta_0)] \quad (15)$$

where $\tau_{muscle}$=10 ms represent the time scale with which a muscle responds to a motor drive and $w_{\Theta}$ is a weight that give the force resulting from a specific drive. The last term on the right-hand side represents a weak decay back to the initial joint position of the limb. Joint angles were limited to be within [0,$\pi$].

#### Data availability statement

Data are available on reasonable request from the corresponding authors.

#### Code availability statement

Code are available on reasonable request from the corresponding authors.

### Supplementary Information

Five experimental data sets that fulfilled the requirements of both successful recording from large numbers of neurons, six motor nerve recordings and activation of distinct motor behaviors was acquired. Summary of the parameters is shown in table 1. The electrode depths are indicated with respect to the ventral side, which puts the electrode arrays in Rexed laminae VII-VIII, where the motor-related inter-neurons are located.

##### Rotational dynamics

To substantiate the observation of rotational population dynamics (Fig. 1), we analysed several trials (n=8) of the same animal in similar manner (**Extended Data Fig. 1a**). The population activity had similar sequential activity. The sorting and PCs are the same as in Fig. 1 and based on trial 3 (indicated by "*"). The distribution of phases, which was calculated with respect to the rhythmic activity of a motor nerve (hip flexor), revealed no clear phase-preference (**Extended Data Fig. 1b**). The population activity in PCA space indicates similar rotational dynamics (**Extended Data Fig 1c**). Analysing all animals in the same manner yielded similar results: Sequential activity in the neuronal population with a continuous cycling through all phases (**Extended Data Fig. 2a**). Hence, most of the phases were represented and there was no distinct mode to see in the histograms (**Extended Data Fig. 2b**). The trajectory in PCA space also had resemblance to rotation in other animals (**Extended Data Fig 2c**). The dynamics was generally low-dimensional, as manifested by most variance was represented by few principal components (**Extended Data Fig. 2d**).

##### Tangling of network and nerve population activity

As established above, the neuronal population trajectory tend to perform a rotation in neural space. Circular motion has the property that trajectories moving in opposite direction is farthest apart and hence they have low tangling (**Extended Data Fig. 3**a). This is in opposition to the traditional half-center model where the neural trajectory would move on a line between two states. Since the trajectories that move in opposite direction are close to each other, such half-center dynamics has high tangling. Hence, the tangling measure ($Q$) can be used to quantify the degree of "alternation" versus "rotation". The nerve output is primarily composed of flexors and extensors, which tend to alternate in opposition. Therefore we quantified the tangling for both the network trajectories, $Q_{network}$, and the associated nerve output trajectories, $Q_{nerve}$, in 3 dimensions. The fraction of observations where $Q_{nerve} > Q_{network}$ was quantified for all trials and data sets and they were all substantially larger than 50% (**Extended Data Fig. 3**b). This suggests that the network is executing a "rotation" whereas the nerves are performing "alternation". This was consistent across the data sets (**Extended Data Fig. 3c-g**).

##### Activity of excitatory and inhibitory neurons in the BSG-model

The Balanced Sequence Generator (BSG) model (Fig. 2) exhibits rotational dynamics, in accord with experimental observations. Here, we further investigate the inhibitory and excitatory population activity during this rotation. In the traditional half-center model, excitatory (E) and inhibitory (I) populations should alternative in activity due to reciprocal connections. In the BSG-model the behavior of the E/I populations is different. Activating the BSG-model with an input drive causes the neurons to oscillate (**Extended Data Fig. 4a-b**). Segregating the population into the E and I and performing the same type of sorting according to phase, there was a similar sequential activity within the E- and I- populations (**Extended Data Fig. 4c-e**). Therefore, looking at these populations individually, they also demonstrate rotational dynamics (not shown). This can be considered an experimental prediction of the BSG-model, that make it distinguishable from the half-center model. Another way of showing these properties is by plotting what is called the eigenmode of the network activity, i.e. the eigenvector corresponding to the largest eigenvalue (**Extended Data Fig. 4f**). Here, each dot represent the activity of a given neuron, in terms if its phase, as the angle in the polar plot, and its firing rate as the radius in the polar plot. The activity is scattered around the origin with many different angles and radii. The different angles indicates that there is no particular phase preference within the population. Similar way of plotting the population activity have been applied in previous reports[26,27]. Segregating the eigenmode into the E- and I-populations, show similar patterns (**Extended Data Fig. 4g-h**). Histograms of the phase of the activity across the neuronal population did illustrate wide phase distribution (**Extended Data Fig. 4i-k**). This should be compared with the experimental observed phase distributions (**Extended Data Figs. 1b and 2b**).

##### Control of force by input drive via the radius of rotation

In the BSG-model the control of amplitude was accomplished by modulating the general neuronal gain. When an increase in gain was provided to the whole network, the dynamics of the network becomes slightly more unstable. This was seen as an expansion of the eigenvalue spectrum of the connectivity matrix (**Extended Data Fig. 5a**). When some of the eigenvalues crossed the stability line (red vertical line) the network started to oscillate. Sorting the neurons according to phase, a sequential activity was revealed (same sorting throughout, **Extended Data Fig. 5b**). As the drive (gain) increased, the amplitude also increased. This was also seen in the PCA where the radius of rotation expanding increasing (**Extended Data Fig. 5c**). Some of the neurons projected to motor neurons resulting in flexor and extensor ENG activity that also increased in amplitude (**Extended Data Fig. 5d**). Note that this control of force did not seem to affect the period of oscillation. This is an advantage of the BSG-model, since other models have not been able to secure independent control of period and amplitude. The correlation between drive, firing rates, nerve output and rotation radius is shown in (**Extended Data Fig. 5e-h**). In the experiments, we observed trials that had different radius of rotation (**Extended Data Fig. 1c**) that was correlated with the ENG amplitude (Fig. 3e-g). This was further analysed by dividing each trial up in smaller pieces and comparing the PCA rotation and nerve activity (**Extended Data Fig. 6a-f**). The nerve output (RMS) had a significant correlation with the radius of PCA rotation both in the presented animal (**Extended Data Fig. 6g**) and across all 5 data sets (**Extended Data Fig. 6h**).

##### Mean and variance of population activity: BSG- model and experiment

In the above we saw that, when the descending input provided activation of the network, the neurons started to oscillate. The descending drive can come from many sources, and in the present experiments the source was cutaneous sensory input that elicit a scratching response. Besides starting the rhythmic activity, the variance in activity was also increased, in both the BSG-model and in the experiment (**Extended Data Fig. 8**). The variance tended to be larger than the excursions in mean firing rate itself. Nevertheless, it is interesting that in spite of the random connectivity the mean value in the model has an oscillatory component, which could explain the sinusoidal cord dorsum potentials, which are known to occur e.g. in cat spinal cords during fictive rhythmic scratching[28,29].

##### Multifunctionalism in the BSG-model and experiment

In figure 5, we observed that both the BSG-model as well as the sampled neuronal population in the experiment, was able to participate and generate two distinct motor behaviors, the pocket- and rostral scratching). Multifunctional circuitry in the spinal cord is well-established[8,30,31], but the mechanisms of how a network accomplishes this is an open question. The same BSG-network was able to generate these two patterns by activation of selected subset of neurons that could shift the phase of one output motor nerve (hip extensor), thus providing two distinct motor programs, pocket- and rostral scratching (Supplementary Video 3). The BSG-nerwork was in fact able to generate many distinct motor patterns, depending on what which gain profile that was activated by descending drive. Here, we explore the diversity in motor output by selecting different random gain profiles to a subset of neurons in the network (**Extended Data Fig. 9a**). This resulted in similar albeit not identical sequential activities (**Extended Data Fig. 9b**), and rotational population dynamics (**Extended Data Fig. 9c**). The resulting motor nerve output of 4 nerves had various and different patterns (**Extended Data Fig. 9d**). The population activity had a more clear and repeatable sequence, when the animal was repeating the same motor output, compared with a different motor behavior. Nevertheless, there was similarity in the sequence of patterns despite the difference in motor programs. The BSG-network seems to have a large degree of flexibility and we suggest that, if properly activated, it could in principle generate any rhythmic motor pattern.

##### Other observations: Non-resetting deletions

A 'deletion' is defined as a brief spontaneous failure of appearance of either a flexor or extensor burst in an otherwise normal robust alternating activity[18,32]. If the sequence of the rhythm starts over after the deletion, it is referred to as a 'resetting deletion'. However, if the rhythm as a whole continues uninterrupted, the deletion is referred to as a 'non-resetting deletion'. The appearance of non-resetting deletions is generally considered evidence for a multi-layered organization within central pattern generators[17–20]. Here, we observed the phenomenon of non-resetting deletions in a subset of trials ($n = 6$) (**Extended Data Fig. 10**). The appearance of such phenomenon is in line with previous observation in this preparation[32], and it suggest to support the notion of multiple layers. Interestingly, the rotational dynamics continued almost unaffected despite the occurrence of a deletion. Nevertheless, in all trials, the part of the neural trajectory where the deletion occurred (red dots, **Extended Data Fig. 10**) was distorted. A distortion of the neural trajectory can provide a simple explanation for the phenomenon, which does not require multiple levels. This suggests that a deletion is a temporary disturbance in the neuronal population dynamics, which is large enough to bring the population trajectory below threshold for eliciting a motor nerve burst for a given muscle.

##### Deletions explained by the BSG-model

If this characterization of the phenomenon of deletions is suitable, it should be a simple task to verify it in the BSG-model. As it turns out, the generation of deletions is straightforward in the BSG-model, where it is not necessary to construct multiple layers in the network. If we imagine the input drive to the network can fluctuate, e.g. due to some biological variability in either the descending drive commands, fatigue or slowing of sensory input, the neural trajectory will get distorted (**Extended Data Fig. 11a-c**). If the drive is low enough (red dots) the neural trajectory will be perturbed below a certain level such that the motor burst will be absent (compare red and blue dots **Extended Data Fig. 11d-g**). Despite the distortion in the trajectory, the rotational dynamics continues to cycle through the phases, hence allowing the next part of the motor sequence to proceed without a reset. In this way, non-resetting deletions can explained by the BSG-model. In fact, these events can be considered a special case of multifunctionalism, perhaps containing an indicating of pathology.

##### Linear decoding of neuronal population activity into nerve output

The transformation of neuronal population activity to motor commands has been investigated in the cortex[14–16]. To our knowledge it has not yet been done in spinal neuronal networks. Since it is important for understanding the issue of deletions, we performed an analysis of the linear transformation of the population activity to nerve output (**Extended Data Fig. 12**). First, a training set containing several trials was used to estimate the linear filter (**Eq. 9**). Once the filter function was obtained, the transformation was validated using a new trial where the predicted nerve output was compared with the actual (**Extended Data Fig. 12b**). The correlation coefficient between predicted and actual was calculated for each nerve and each trial. This was also done for two motor behaviors (left and right pocket scratching). There was a mean level of 0.6 in correlation coefficient in both behaviors (**Extended Data Fig. 12c**). Similar analysis was performed for the data containing deletions (**Extended Data Fig.**

13). The occurrence of deletions seems to be somewhat predicted by the linear decoder (compare red arrows **Extended Data Fig. 13b**).

##### Overlap in subspace representation of one behavior compared with another

The principal components for the network and nerve activity was first calculated for one behavior and one trial (left pocket scratch). Then the activity of a different trials of same behavior as well as trials of a different behavior (right pocket scratching) was projected onto these principal components. The variance captured by these projections was then calculated for the first three components and compared with the variance that could have been captured if the principal components of the given trial itself, i.e. in its "native" space. The overall observation is that trials of the same behavior has larger overlap ($\sim$0.7) compared with the projections from different behaviors (cf. green and orange **Extended Data Fig. 7a**). Second, the subspace overlap in the nerve activity was generally larger than that for the network, presumably because the nerve space has only 6 dimensions (**Extended Data Fig. 7b**). There was a significant almost linear relationship between the subspace overlap in the nerve activity and that of the network (**Extended Data Fig. 7c**), which is an indication that when the motor nerve patterns are more similar then the network population activity is also more similar.

**Table 1. Overview of data**. Units column represents the number of neurons (units) that was isolated using polytrode spike sorting. Number of trials within each behavior (ipsilateral or contralateral pocket scratching or rostral scratching). The vertebral location indicate where the Berg64-electrode probes were inserted. The VD (ventrodorsal) depth indicated how deep the probes were inserted from the ventral side (preparation was upside-down).

| Data set | Units | Behaviors | Trials (ipsi-,contra-pocket, rostral) | Vertebral location | VD-Depth ($\mu m$) |
|---|---|---|---|---|---|
| 1 | 226 | 3 | 14 (6/6/2) | D8, D9, D10 | 560, 560, 750 |
| 2 | 249 | 3 | 10 (4/3/3) | D8, D9, D10 | 1050, 1235, 1050 |
| 3 | 214 | 2 | 19 (10/9/0) | D8, D9, D10 | 702, 694, 725 |
| 4 | 58 | 3 | 16 (5/5/6) | D8, D9, D10 | 1050, 1060, 1500 |
| 5 | 200 | 3 | 23 (6/6/11) | D8, D9 (contralateral), D10 | 400, 400, 400 |

### Mathematical note

In the following we outline how the generation of network oscillations in our model can be understood in the general framework of linear dynamical systems, and how the oscillation depends on the eigenvalue spectrum of the effective network connectivity.

In general the activity $\mathbf{r}$ of a dynamical system can be described by a set of ordinary differential equations written in compact notation as:

$$\frac{d}{dt}\mathbf{r}(t) = f(\mathbf{r}) \tag{16}$$

where function $f$ describes both network interactions as well as intrinsic properties of the neurons in the network.

Let us start with a simple network model where $\mathbf{r}(t)$ represents a vector containing the firing rates (relative to a baseline) of $N$ leaky neurons and the matrix $\mathbf{W}$ describes that synaptic network interactions around the baseline rates:

$$\tau\frac{d}{dt}\mathbf{r}(t) = -\mathbf{r} + g\mathbf{W}\mathbf{r} \tag{17}$$

Here $\tau$ is the time constant for change in the firing rate and $g$ is the slope (gain) of the firing rate function that we assume is linear. This network has an equilibrium point, i.e. a point where $\frac{d}{dt}\mathbf{r}(t) = 0$, at $\mathbf{r}(t) = 0$, corresponding to state in which all neurons fire at the baseline rate. After rewriting Eq.17 slightly to

$$\frac{d}{dt}\mathbf{r}(t) = \frac{1}{\tau}[g\mathbf{W} - \mathbf{I}] \cdot \mathbf{r} \tag{18}$$

and defining $\mathbf{A} = \frac{1}{\tau}[g\mathbf{W} - \mathbf{I}]$ we see that our network has the general form of a linear dynamical system

$$\frac{d}{dt}\mathbf{r}(t) = \mathbf{A} \cdot \mathbf{r} \tag{19}$$

If an input (perturbation) to the network at $t = 0$ is in the direction of one of the eigenvectors $\mathbf{x}_k$ of $\mathbf{A}$ (i.e. population vectors for which $\mathbf{A}\mathbf{x}_k = \lambda_k \mathbf{x}_k$, where $\lambda_k$ is the corresponding eigenvalue) the dynamics is given by $\frac{d}{dt}\mathbf{r}(t) = \mathbf{A} \cdot \mathbf{x}_k = \lambda_k \mathbf{x}_k$, which has the solution

$$\mathbf{r}(t) = \mathbf{x}_k e^{\lambda_k t} \tag{20}$$

Since any perturbation can be expressed as a sum of eigenvectors the stability of the equilibrium point is therefore determined by the distribution of eigenvalues $\lambda_k$ of the matrix $\mathbf{A}$.

If all eigenvalues have a real part smaller than zero the activity $\mathbf{r}(t)$ decays back to the equilibrium. In contrast, if at least one eigenvalue has a real part greater than zero the activity of the network is unstable and will grow exponentially due to Eq.20. The crucial parameter for the stability of the network is thus the largest real part of all the eigenvalues: $\Lambda = \max_k Re(\lambda_k)$.

Let us next assume that the connectivity matrix $\mathbf{W}$ of our simple network model is balanced, i.e. that excitatory and inhibitory inputs are equally strong for every neuron in the network. Then the eigenvalues spectrum of the static connectivity matrix $\mathbf{W}$ is distributed on a disc in the complex plane centered at zero with a spectral radius determined by the variance of the connectivity[25].

To further analyze our simple network model it is of interest to find the eigenvalues of the matrix $\mathbf{A} = \frac{1}{\tau}[g\mathbf{W} - \mathbf{I}]$. Since the subtraction by the identify matrix $\mathbf{I}$ only shifts the eigenvalue spectrum by -1, the stability criterion outlined above for the largest eigenvalue of $\mathbf{A}$ is equivalent to the largest eigenvalue of the effective connectivity $\mathbf{W}_{\text{eff}} = g\mathbf{W}$ having a real part that crosses 1. Adjusting the gain $g$ of the firing rate function adjusts the spectral radius of $\mathbf{W}_{\text{eff}}$ so that the gain can be used as a parameter to adjust the stability of the network.

The eigenvalues of a specific realization of $\mathbf{W}$ may result either in a scenario where the imaginary part of $\lambda_{\max}$ is either zero or non-zero. If the imaginary part is non-zero the solution to Eq.20 is oscillatory while if the imaginary part is zero the solution is non-oscillatory.

### Supplementary videos - captions

**Supplementary Video 1: Rhythmic motor activity and neuronal population dynamics in the spinal cord.** The first sequence shows the nerve activity (top, left), the raster plot of the neuronal population (middle, left), and the estimated firing rates (bottom, left) during a motor program (pocket scratching). The first two principal components are shown (right). The sound is the occurrence of spiking across the whole population. The next sequence is the first three principal components of the population firing. Sound is the occurrences of spiking across the whole population, which sounds different because time has been slowed down. Trial 3, data set 3. The last sequence shows the evolution of the first three principal components of 10 trials together in various colors. Data set 3.

**Supplementary Video 2: Comparison between experiment and the BSG-model.** Left: Experiment, Right: BSG-model. Four motor nerves (top). Sorted firing rates for the neuronal population (Middle). First two principal components (bottom). Network is activated by a constant increase in input. Experiment: Trial 1, data set 3.

**Supplementary Video 3: Transition between two behaviors of the BSG-model by selective activation.** A certain mode of activity within the network was generated by a distinct gain-modulatory input to some neurons ("gain-profile") in the BSG-network (top, left). This entailed rotational dynamics, as shown in PCA (middle, top) and the sequential firing rate activity (middle). The nerve output to the knee flexor and extensor, and hip flexor and extensor (bottom) followed a pattern that resulted in one distinct motor behavior of the hind limb (pocket scratching, top right). Halfway through the sequence (red line), the mode of activity was switched by changing the input pattern (top, left). The network still had rotational dynamics, but with a slight modification (Compare orange and blue lines, PCA and limb trajectory). The new network input caused a different motor behavior (behavior 2), which was rostral scratching. The phase of the muscle contraction had been changed as seen in the nerves (bottom). In spite of the difference in the motor output the rotational dynamics only had minor changes (PCA) and the neuronal sequence of activity was similar (sorted firing rates).

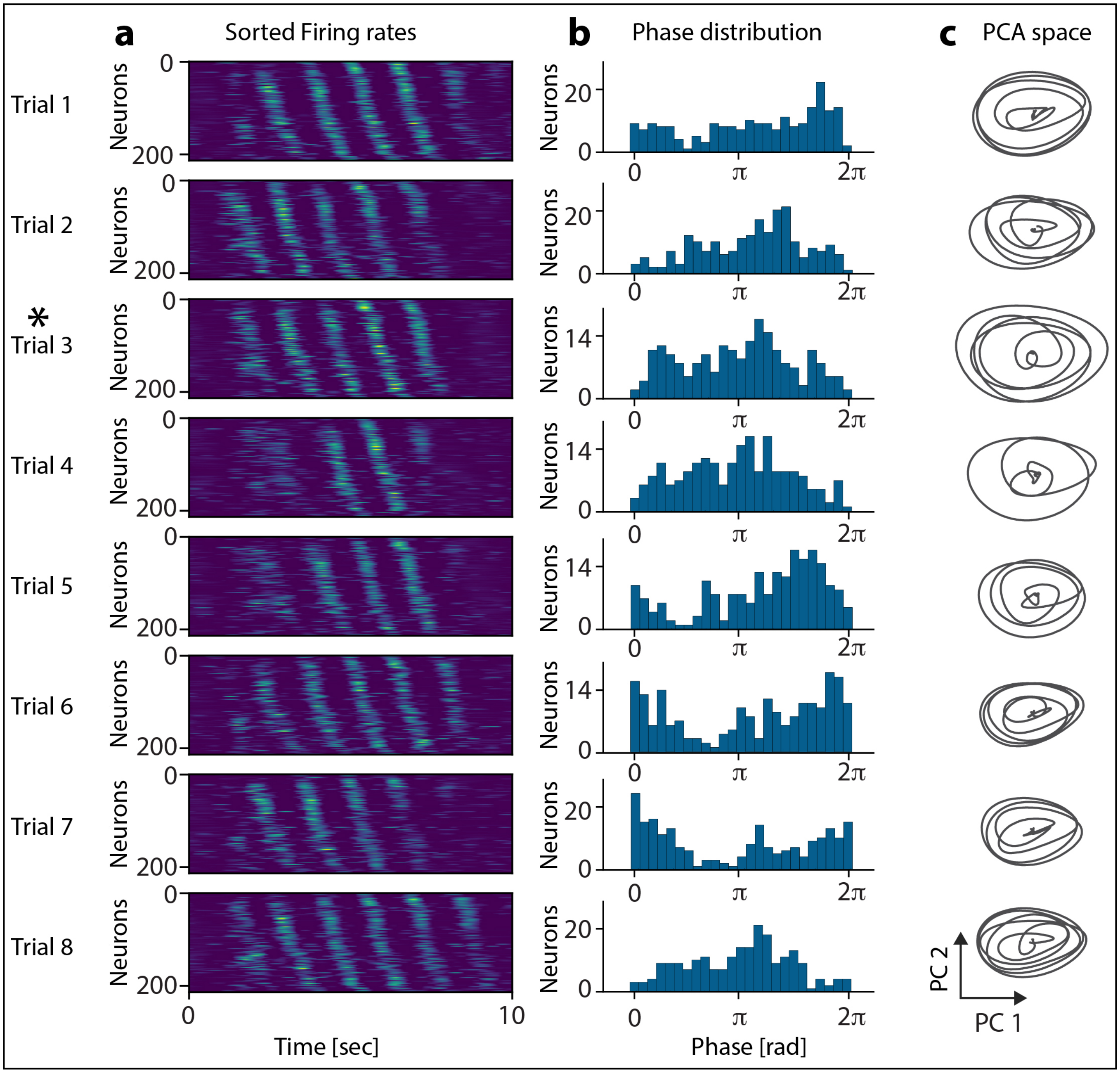


**Extended Data Fig. 1.** | **Rotational population dynamics across trials in the lumbar spinal motor network during rhythmic movement**. **a**, The firing rates (normalized, color coded) of 214 spinal neurons in laminae VII-VIII as a function of time and sorted according to phase with respect to the nerve activity (hip flexor). Eight consecutive trials from same experiment with a 5 min pause in between each. **b**, The phase distribution across the neuronal population. **c**, The population activity has rotational dynamics, as demonstrated by the circular motion of the first two principal components. The PCs were calculated by the data of one trial (trial 3, "*") and the applied to the rest of the trials. The sorting of neurons was according to their phase relation with representative nerve for one trial (also trial 3, "*") and this order was maintained for the rest of the trials. Bottom scale bars represent 1000.

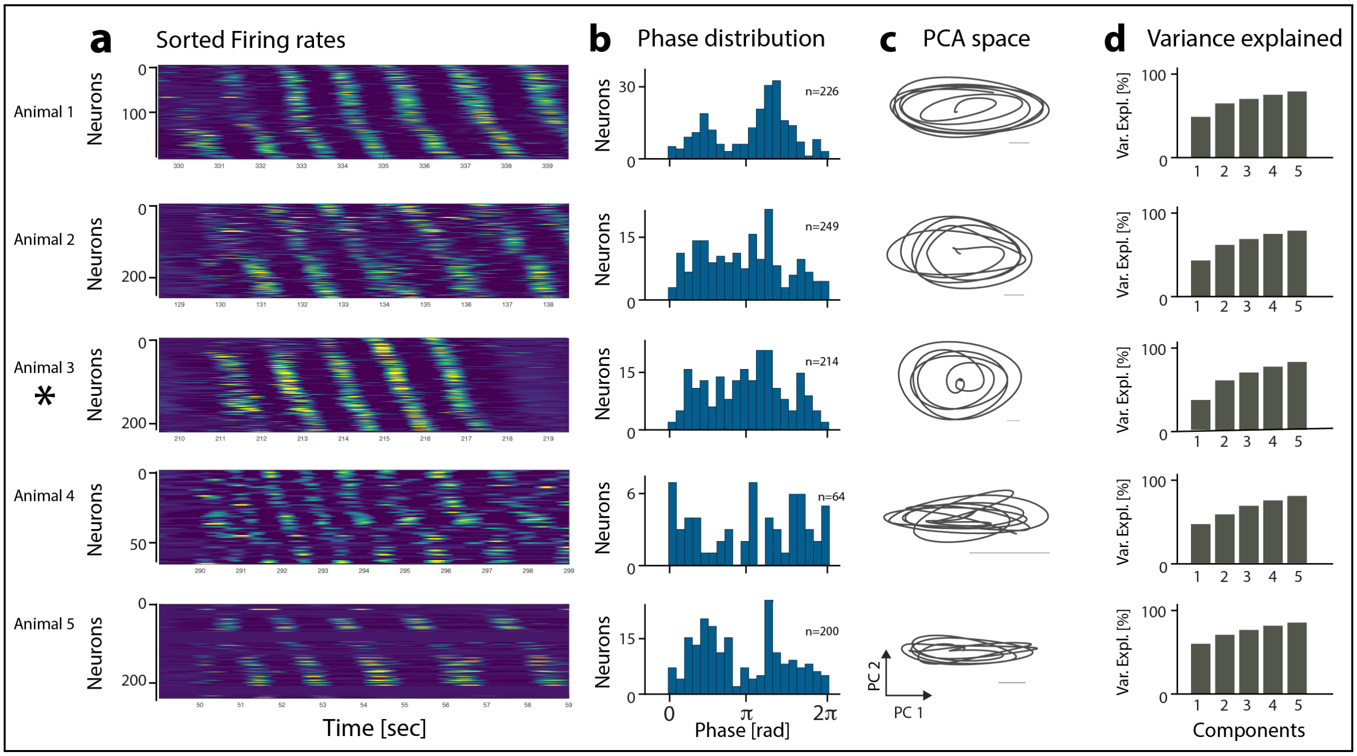


**Extended Data Fig. 2.** | **Rotational population dynamics in a spinal motor networks across animals. a**, The rhythmic firing rates in populations of spinal neurons in laminae VII-VIII shown in colors as a function of time and sorted according to phase with respect to a nerve (hip flexor). A representative trial from 5 experiments of approximately 10 seconds demonstrate similar sequential/rotational population activity. Animal used in Extended Data Fig. 1 is marked "*". **b**, The corresponding distribution of neurons having preferred phases among the population of rhythmic neurons. **c**, Population activity represented by first two principal components exhibit rotational dynamics. Scale bars: 250. **d**, Cumulative explained variance by principal components, indicating the population dynamics is low-dimensional, i.e. most of the variance is captured by few components.

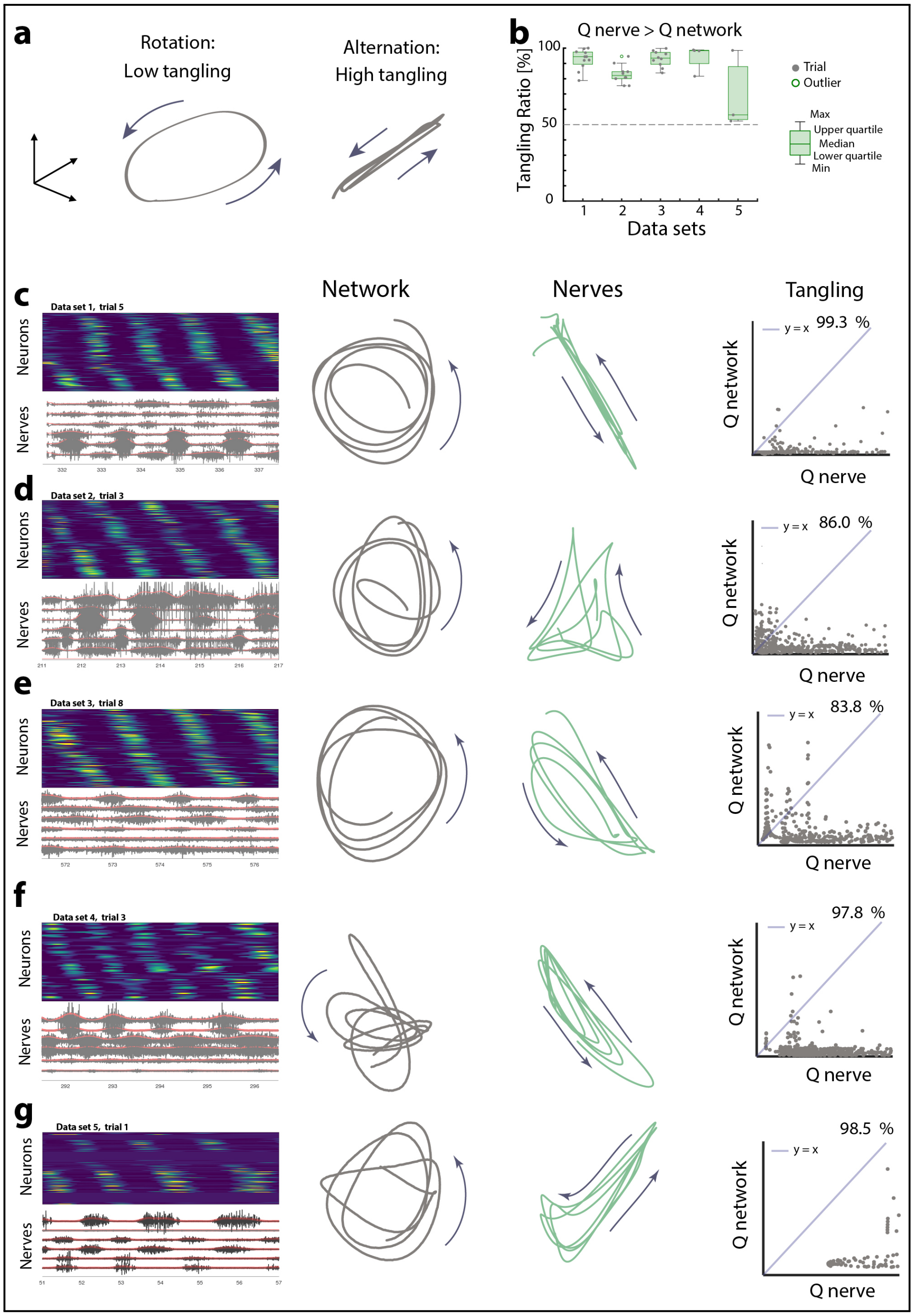


**Extended Data Fig. 3.** | **Neuronal population trajectories in PC-space has lower tangling than the corresponding motor nerve trajectories. a**, Illustration that during rotational dynamics the points in the trajectory that move in opposite direction are also far apart, i.e. they have low tangling (left), whereas during alternation the points of the trajectory that move in opposite direction are also close, i.e. have high tangling (right). **b**, The ratio of tangling metric[21] of the principal components (PCs) trajectory of the nerves ($Q_{nerve}$) to that of the network ($Q_{network}$). This ratio is close to 100%, which indicates most trials and animals had a larger tangling of the motor nerves than the network. **c-g**, Sample trials from 5 different data sets. Left is shown the phase sorted firing rate activity (top) and the associated nerves (bottom). The nerves were rectified and low-pass filtered (red) on temporal scale matching the firing rates. The PCs of network (middle left) and nerves (middle right, green). Scales of PCs are variance-normalized. The tangling metric ($Q$) for the nerve PCs (in 3 dimensions) is calculated as a function of time ($t$) through the trial and plotted versus that for the network. The ratio of points below the $x = y$–line (pale blue) is indicated in percent and form one point in panel (b). Note that the nerve trajectories more resembles "alternation" whereas the network more resembles "rotation"-scheme of (a).

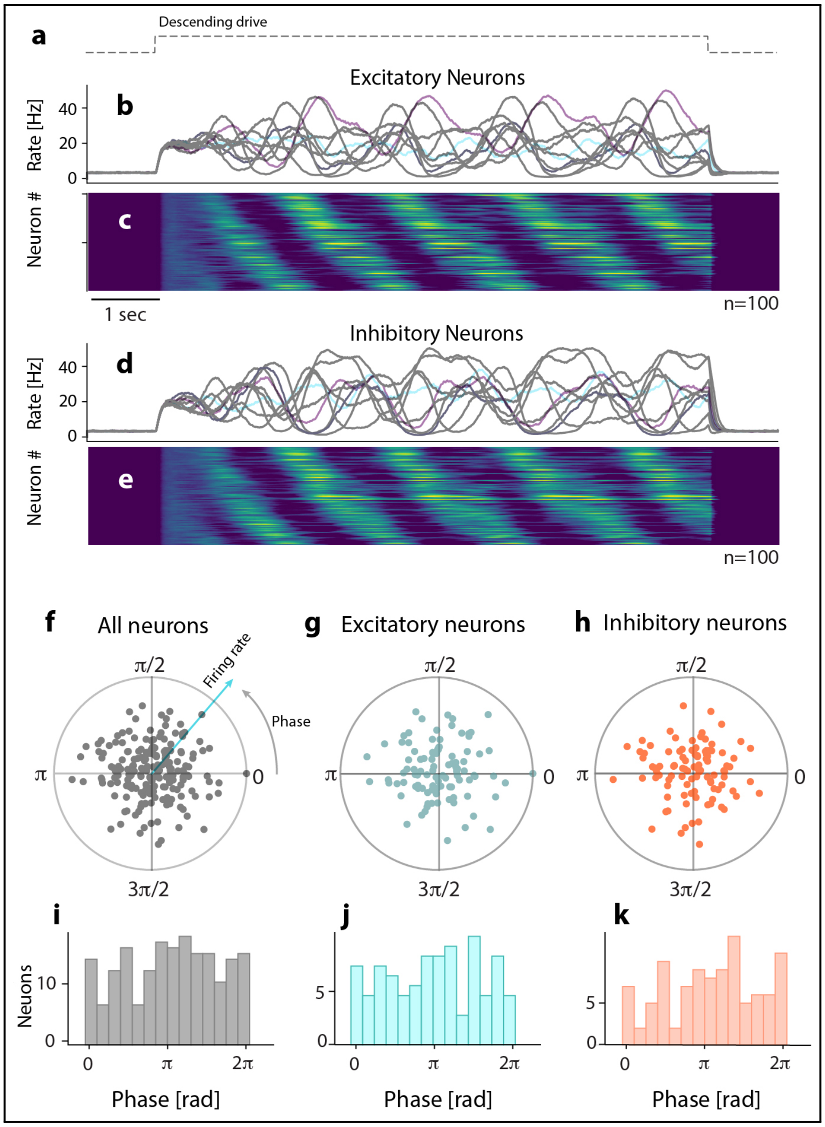


**Extended Data Fig. 4.** | **Rotational ensemble activity within the excitatory and inhibitory sub-populations in the BSG-model. a**, Activation of the motor circuit by descending drive. **b**, The firing rates of 10 sample excitatory neurons as a consequence of the descending input. **(c)** Sorting the excitatory neurons according to phase of firing rates reveals a sequential activity similar to the previously observed for all neurons. **d-e**, Activity and similar sorting of the inhibitory sub-populations reveals similar sequential and rotational dynamics within that sub-population. **f**, The network eigenmode for the whole network: Each dot represent both the phase (the polar angle) and the peak firing rate (the radius) for a given neuron (n=200). **g-h**, Similar plot for the excitatory and inhibitory populations **i-k**, the distribution of phases in linear histograms for all neurons (i), excitatory (j) and inhibitory neurons (k). To be compared with experimental distributions (Ext. Data Figs. 1b and 2b).

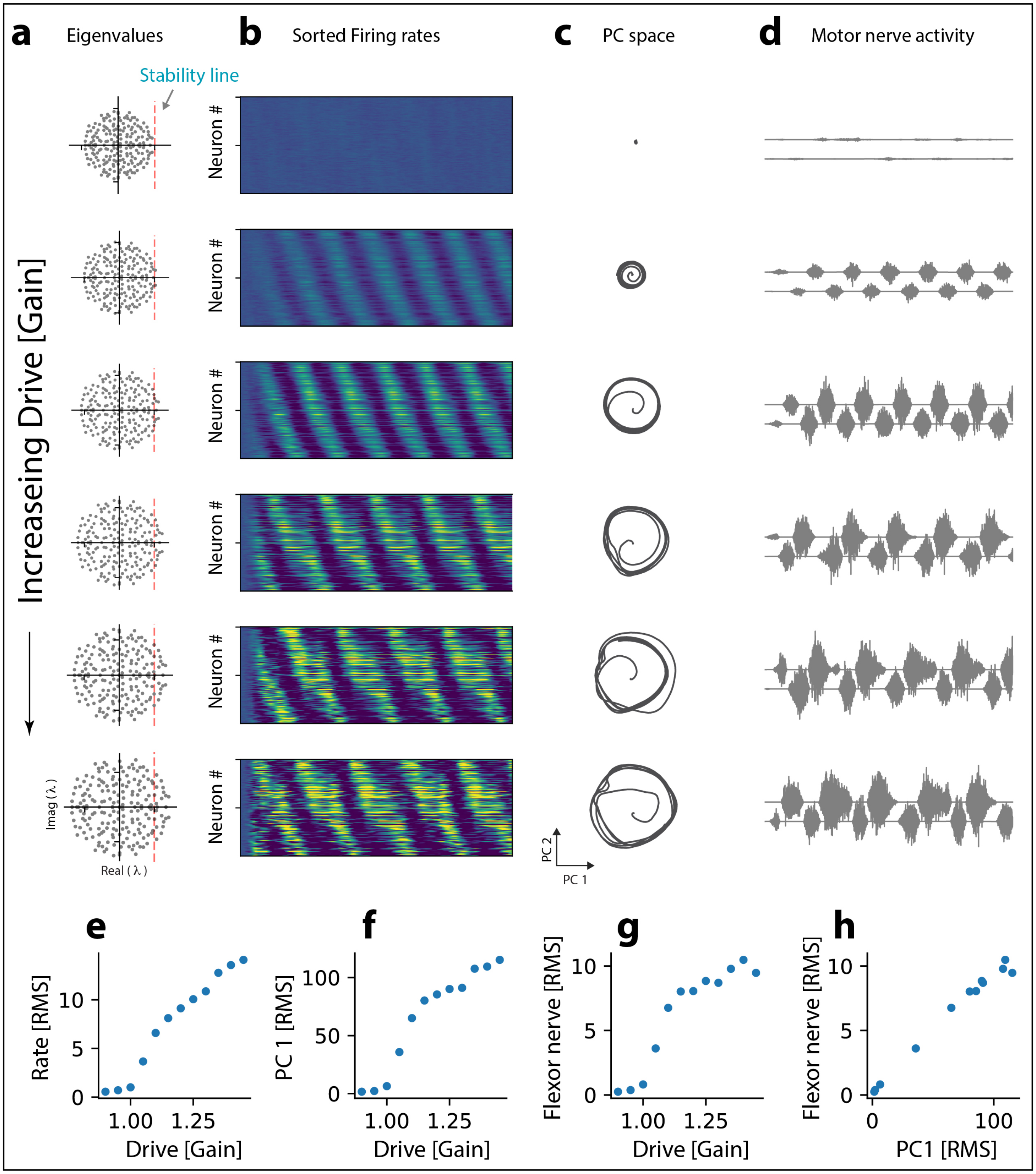


**Extended Data Fig. 5.** | **BSG-model: Correlation between descending drive and radius of rotation as well as amplitude of nerve output without affecting the period.** **a**, For low neuronal gain (top), the eigenvalue spectrum does not have any eigenvalues that cross the stability line (broken vertical line). As the gain increases (downward direction) the spectrum expands and eigenvalues cross the stability line. For larger gain the eigenvalues cross the stability line farther. **b**, The associated population dynamics (sorted firing rates) exhibit oscillation of increasing magnitude as the drive increase. **c**, The rotational dynamics also has a radius that increases with increasing drive. **d**, The resulting motor nerve output is also increasing in amplitude. **e**, Descending drive (gain) versus the population firing rate (RMS), radius of rotation in PC space, **f**, and amplitude of nerve output (flexor RMS), **g**. The radius of rotation (PC1 RMS) vs. the nerve amplitude (flexor RMS).

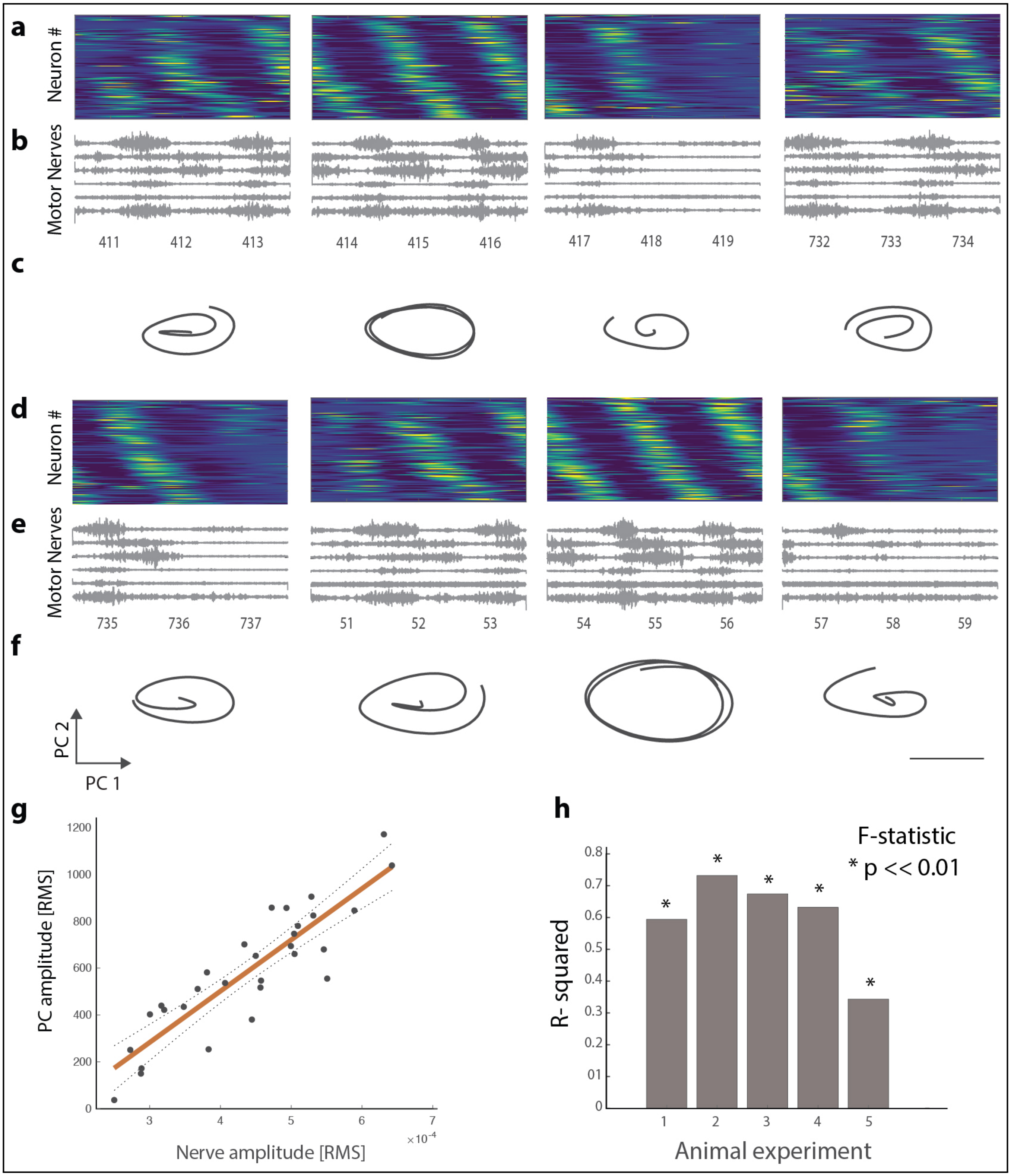


**Extended Data Fig. 6.** | **Radius of rotation correlates with nerve output in experiment. a**, Sample trial where the population activity was divided up in pieces with the corresponding nerve output **b**. **c**, The PC manifolds had rotation with varying radius. **d-f**, other pieces with same organization. **g**, The RMS of the nerve activity versus the RMS of the two first principal components for various pieces of activity had a significant correlation. **h**, The $R^2$ values for all animal tested (n=5). *: F-statistic of rejection of no trend at p«0.01. **f**, Scale bar: 1000.

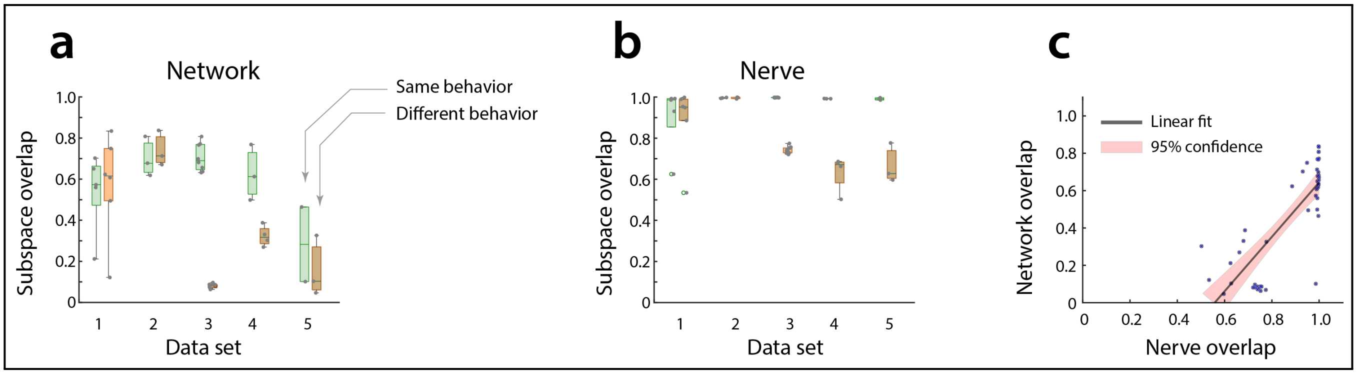


**Extended Data Fig. 7.** | **Representation of one behavior in the subspace of another behavior. a**, The variance captured by the projection of the network dynamics onto the first 3 principal components of another trial (green) normalized by the variance captured by the PCs of its own dynamics. Orange: the subspace overlap of a different behavior. **b**, the subspace representation of the nerve activity of same behavior (green) and a different behavior (orange). **c**, Nerve overlap plotted agains the network overlap. A large overlap in nerve output is associated with a large overlap in network overlap.

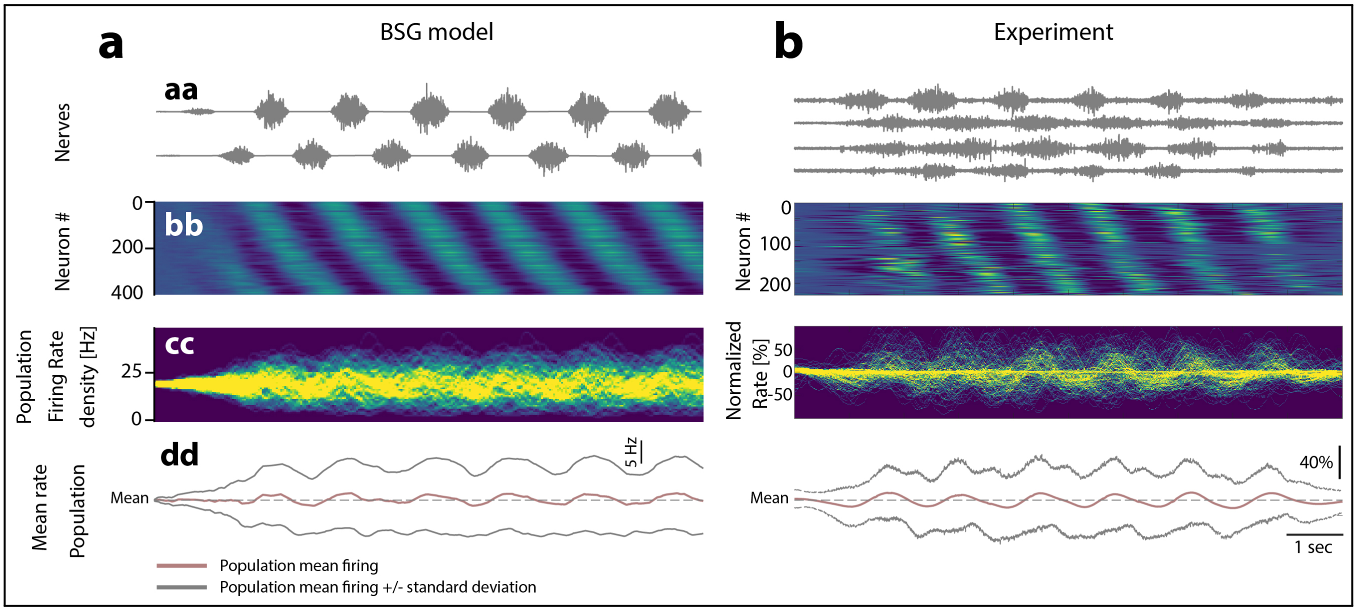


**Extended Data Fig. 8.** | **Variance of population firing rate increase during network output: model and experiment.** **a-aa**, The flexor/extensor nerve output from the BSG-network. **b**, The sorted neuronal population firing rate (n=400 neurons) with rotational dynamics. (cc) Color map of the population firing rate. dd, Mean (red) and variance of the population activity. b same organization as in (a), but for experimental data. Animal no. 3 trial 8.

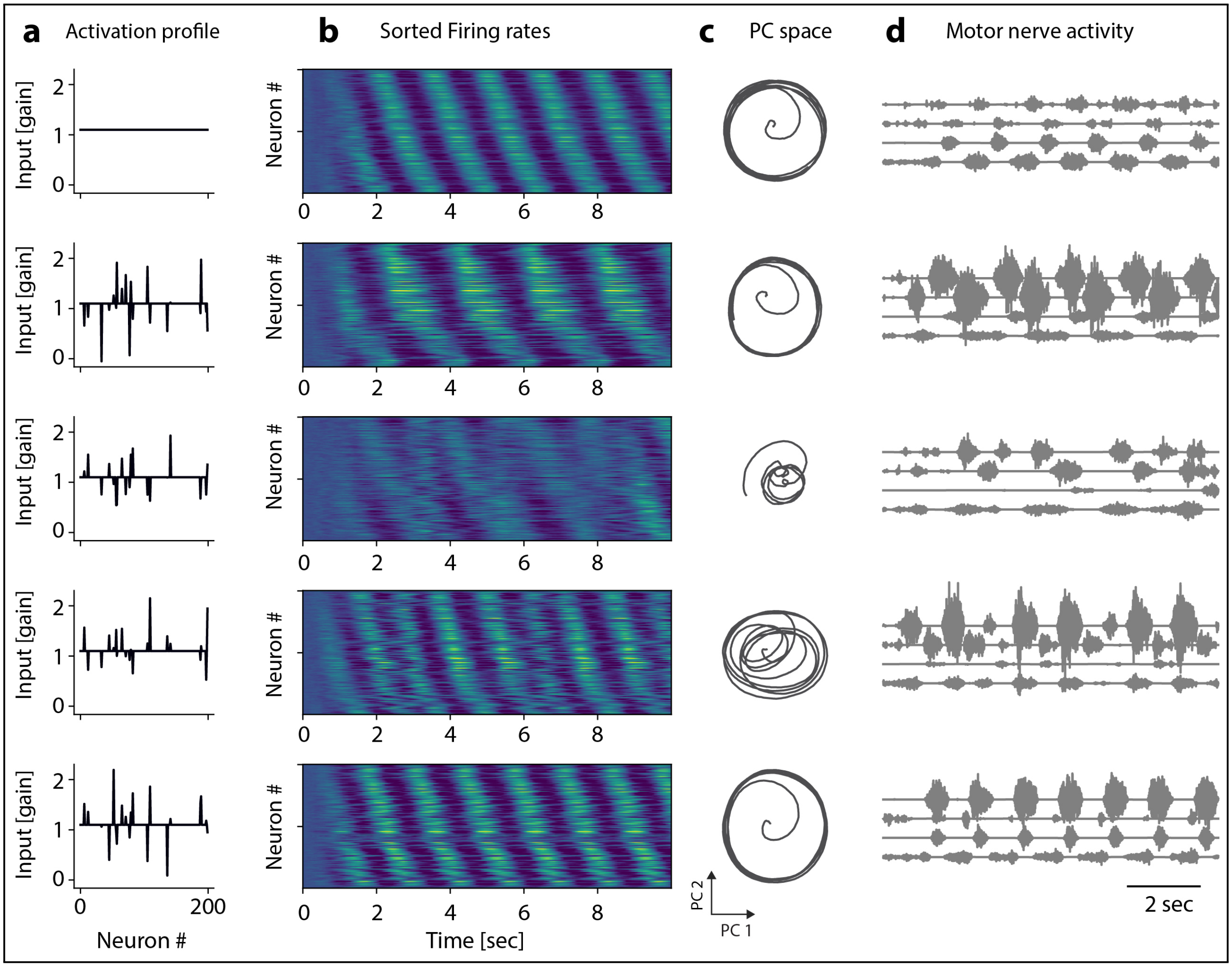


**Extended Data Fig. 9.** | **Multifunctionalism in the BSG-model.** (a) Five examples of specific activation/modulation of selected neurons in the network ("activation profiles"). The top profile has a an even distribution, whereas all the below profiles has selective modulation of specific neurons. (b) The ensemble activity as a result of the activation profile show a sequential activity, with similar but not identical sequence of activity. (c) The first two principal components, based on the top activation profile, all exhibit rotational dynamics, albeit with different radius and trajectories. (d) the output motor patterns associated with the different activation profiles and ensemble activities.

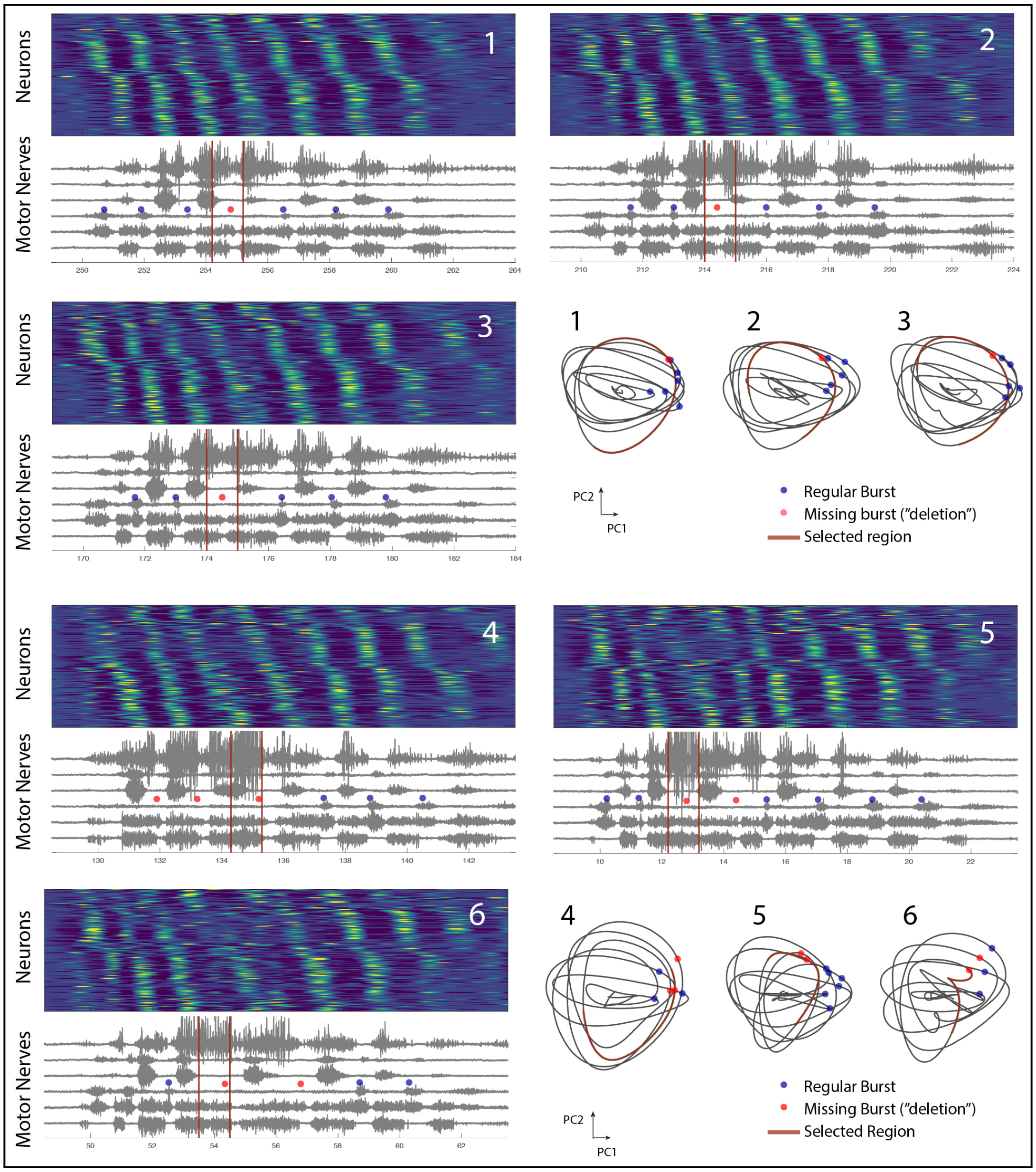


**Extended Data Fig. 10.** | **Trials containing "deletions" have persistent rotational dynamics albeit with distortion in the neural trajectory.** Six trials (1-6) shown with the neuronal firing rates sorted according to phase (color map, top) and the 6 nerves (bottom) during a motor behavior (pocket scratching). The absence of a burst, i.e. a deletion, was observed in the hip extensor nerve recording (red dots) whereas the hip flexor (bottom trace) seems to continue and combine two cycles although with a small decrease. Regular bursts are also indicated (blue dots). The corresponding trajectories in PC-space are shown with the corresponding dots matching the time in the nerve activity. A selected period around the occurrence of one delete is indicated in the nerve traces (red vertical lines). The corresponding time in the trajectory is also indicated in red. Note that deletions tend to occur at smaller radius of the rotation and the population firing rates (color map) are more dim at those instances.

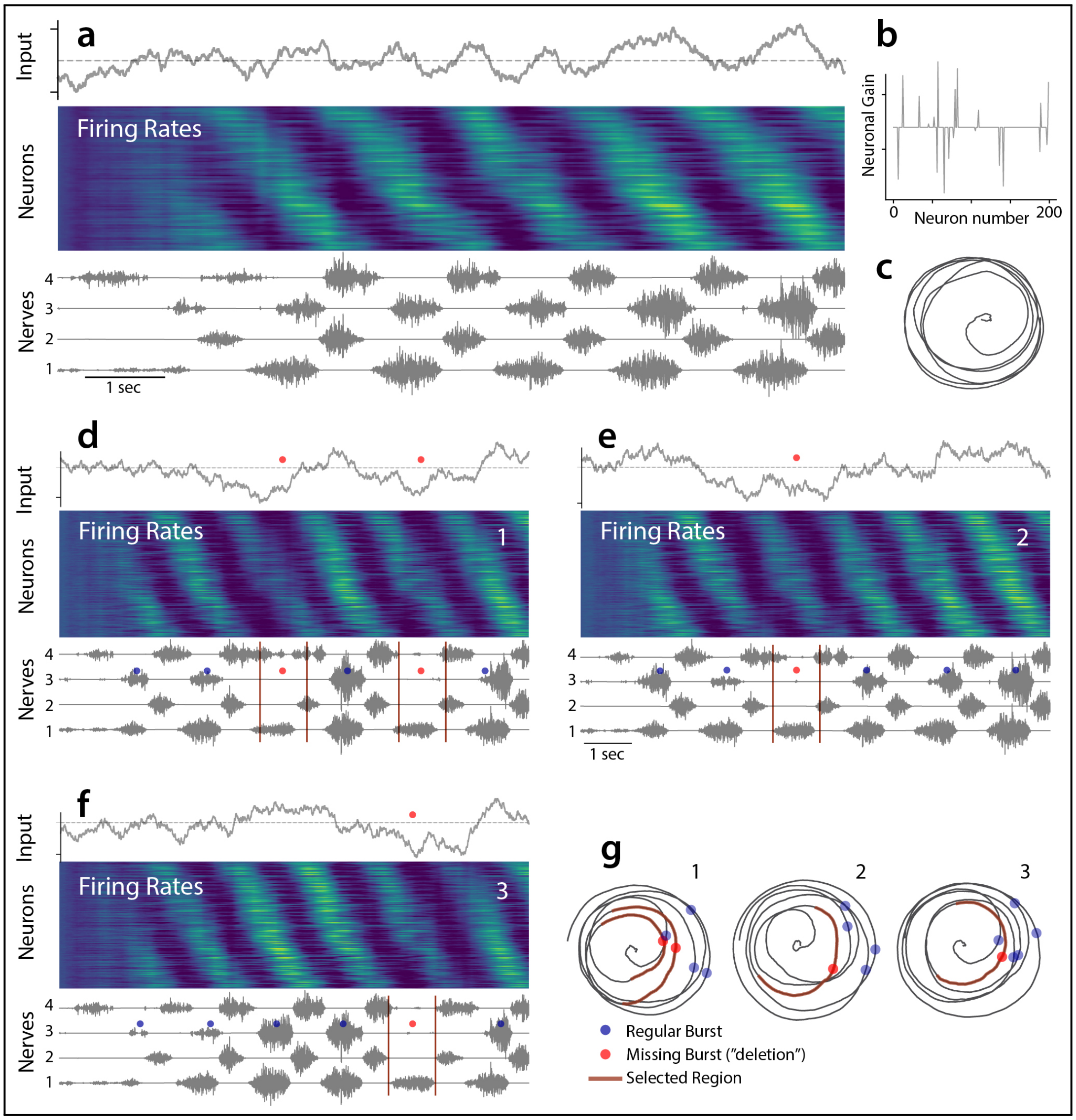


**Extended Data Fig. 11.** | **The BSG-model can explain "deletions" by a varying network input. a**, A proper motor behavior devoid of "deletions" can be produced by the balanced sequence generator despite receiving a varying input (top). The firing rates for the sorted neuronal population (middle), and the resulting motor nerve output pattern (bottom). **b**, The appropriate motor program shown in (a) is achieved by a selective gain-modulation, i.e. gain-profile (y-axis), across the neuronal population (x-axis). **c**, Population activity from (a) represented by in PC-space by the first two components. **d-f**, When the varying input transiently becomes too low at a certain phase the nerve cycle is absent, i.e. a "deletion" has occurred (red dots). The firing rates of the neuronal population will be lower at these instances and hence appear more dim in the color map (middle). A consequent absence of a burst in the nerve is seen (nerve 3, compare red and blue dots). **g**, the PC-trajectories corresponding to (d-f) indicated as 1,2 and 3. The temporary distortion of the trajectory at a particular phase is associated with a deletion (red dots). Red parts of the trajectory represents the period between vertical red bars indicated in the nerve activities (d-f). Compare with Extended data figure 10.

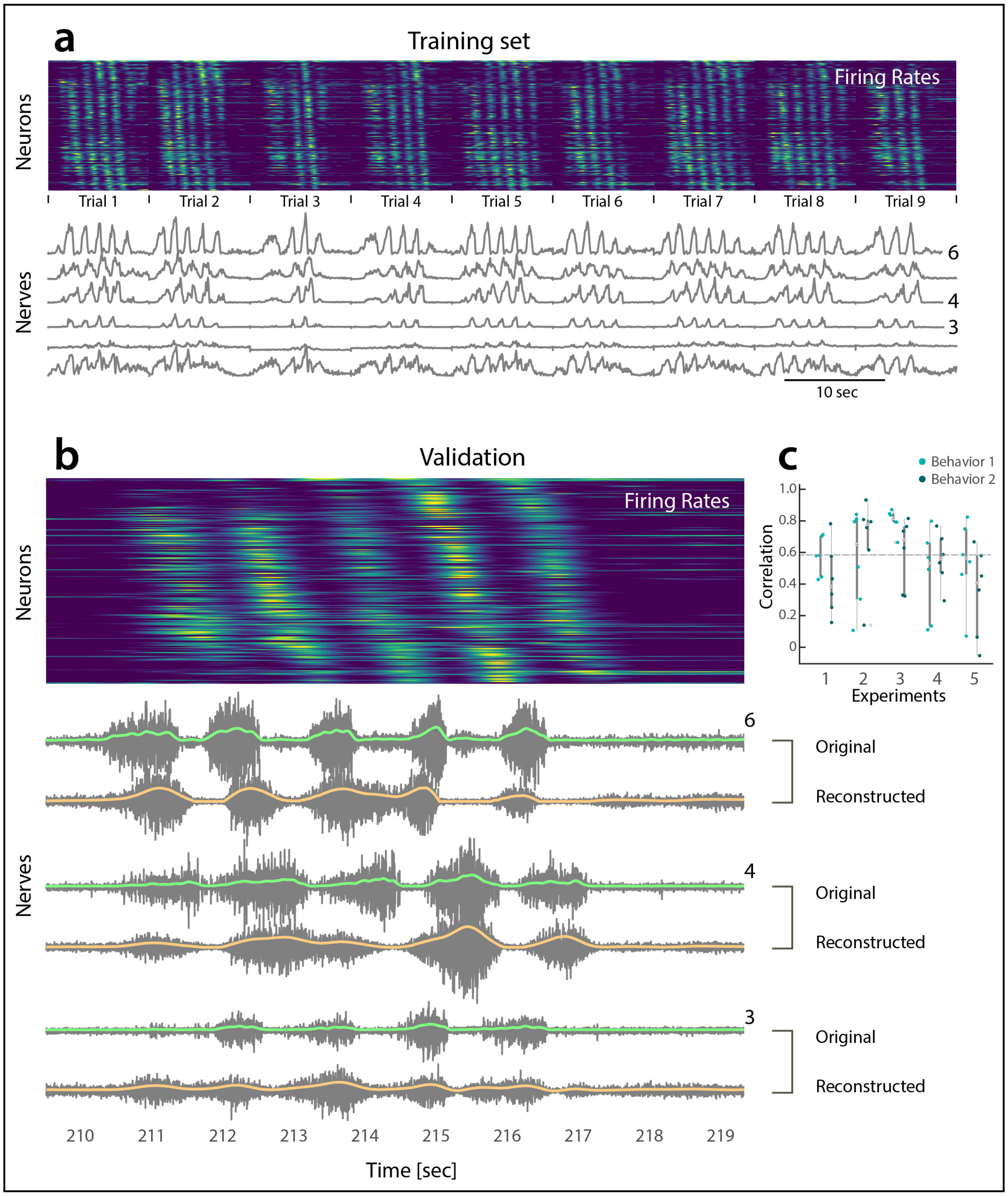


**Extended Data Fig. 12.** | **Reconstruction of nerve output based on linear decoding of neuronal population activity. a**, A linear decoder function was estimated using a training set consisting of 9 trials of same behavior. Top: color coded firing rates for the neuronal population (sorted according to phase) with 9 trials concatenated. Bottom: the rectified and low-pass filtered motor nerve output of 6 nerves. **b**, A trial, that was not included in the training set, is used for validation of the linear decoder. Top: the firing rates of the population, similar to (a). Bottom: The nerve output of 3 selected nerves (rectified and LP-filtered in green, nerves 3, 4 and 6). The reconstructed standard deviations of the nerves (orange) are multiplied by white Gaussian noise to imitate nerve output (gray). **c**, the correlation between predicted and actual nerve output for the six nerves (individual dots) are shown for two different motor behaviors (right and left pocket scratching) across the 5 experiments. The median value across all nerves and experiments is $R = 0.6$. All correlations were significantly different than zero ($p << 0.01$).

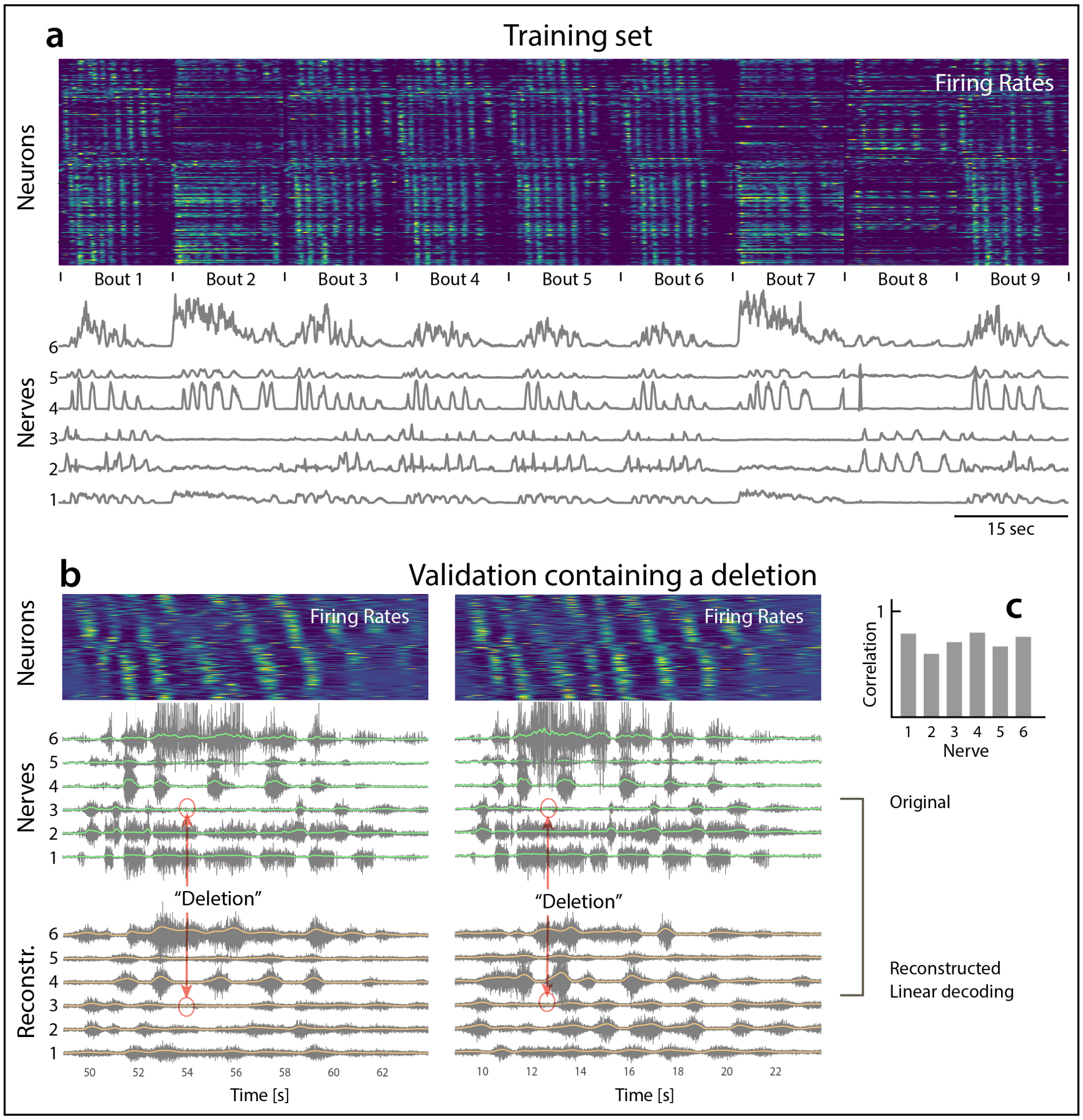


**Extended Data Fig. 13.** | **Prediction of a deletion using a linear decoder. a**, Training set consisting of 9 bouts, ie. trials of different motor behaviors, which is used to train a linear decoder function. Top: color coded firing rates for the neuronal population (sorted according to phase) with 9 concatenated bouts. Bottom: the rectified and low-pass filtered motor nerve output of 6 nerves. **b**, two trials, that was not included in the training set, contained instances of "deletions", ie. . Top: the firing rates of the population, similar to (a). Bottom: The nerve output of 3 selected nerves (rectified and LP-filtered in green, nerves 3, 4 and 6). The reconstructed standard deviations of the nerves (orange) are multiplied by white Gaussian noise to imitate nerve output (gray). **c**, the correlation between predicted and actual nerve output for the six nerves (individual dots) are shown for two different motor behaviors (right and left pocket scratching) across the 5 experiments. The median value across all nerves and experiments is $R = 0.6$. All correlations were significantly different than zero ($p << 0.01$).
